# The Drosophila ZAD zinc finger protein Kipferl guides Rhino to piRNA clusters

**DOI:** 10.1101/2022.05.09.491178

**Authors:** Lisa Baumgartner, Dominik Handler, Sebastian Platzer, Peter Duchek, Julius Brennecke

## Abstract

RNA interference systems depend on the synthesis of small RNA precursors whose sequences define the target spectrum of these silencing pathways. The *Drosophila* Heterochromatin Protein 1 (HP1) variant Rhino permits transcription of PIWI-interacting RNA (piRNA) precursors within transposon-rich heterochromatic loci in germline cells. Current models propose that Rhino’s specific chromatin occupancy at piRNA source loci is determined by histone marks and maternally inherited piRNAs, but also imply the existence of other, undiscovered specificity cues. Here, we identify a member of the diverse family of zinc finger associated domain (ZAD)-C_2_H_2_ proteins, Kipferl, as critical Rhino cofactor in ovaries. By binding to guanosine-rich DNA motifs and interacting with the Rhino chromodomain, Kipferl recruits Rhino to specific loci and stabilizes it on chromatin. In *kipferl* mutant flies, Rhino is lost from most of its target chromatin loci and instead accumulates on pericentromeric satellite arrays, resulting in decreased levels of transposon targeting piRNAs and impaired fertility. Our findings reveal that DNA sequence, in addition to the H3K9me3 mark, determines the identity of piRNA source loci and provide insight into how Rhino might be caught in the crossfire of genetic conflicts.

## INTRODUCTION

Eukaryotic genomes are littered with transposable elements. By constantly aiming to increase their copy number, transposons pose a major threat to genomic integrity and must therefore be tightly controlled. An important transposon silencing mechanism in animals is the PIWI-interacting small RNA (piRNA) pathway (Siomi et al., 2011, Czech et al., 2018, Ozata et al., 2018, Senti and Brennecke, 2010). Its loss leads to transposon de-repression and mobilization, DNA damage, and sterility. At the center of the piRNA pathway are Argonaute proteins of the PIWI clade loaded with 23-30nt long piRNAs which enable sequence specific repression of transposons at the transcriptional and post-transcriptional levels. To control the highly diverse transposon sequence repertoire, an equally diverse set of piRNAs must be generated. Transposon targeting piRNAs are typically encoded in specialized genomic regions (Brennecke et al., 2007, Houwing et al., 2007, Aravin et al., 2008). These so-called piRNA clusters are enriched in transposon sequences, range from a few to several hundred kilobases in length, and act as heritable and adaptive transposon sequence libraries for the piRNA pathway.

Through their transcription, piRNA clusters provide the essential small RNA precursors with sequence information antisense to active transposon transcripts. The definition of specific genomic loci as piRNA precursor sources is therefore of central importance to determine the pathway’s target spectrum. In *Drosophila* gonads, two general types of piRNA clusters are distinguished based on their mode of transcription: Uni-strand clusters are transcribed on one genomic strand from canonical RNA polymerase II promoters, giving rise to long, single stranded, and poly-adenylated transcripts (Brennecke et al., 2007, Mohn et al., 2014, Goriaux et al., 2014). The second type are dual-strand piRNA clusters, which are transcribed on both genomic strands (Brennecke et al., 2007, Mohn et al., 2014, Klattenhoff et al., 2009, Zhang et al., 2014b, Chen et al., 2016, Andersen et al., 2017). Dual-strand clusters are active in germline cells and are embedded within heterochromatin. They differ from canonical RNA polymerase II transcription units by a lack of defined promoters, and by suppression of splicing and cleavage- and polyadenylation signals.

The molecular identity of dual-strand piRNA clusters is conferred by Rhino, a germline specific variant of the canonical Heterochromatin Protein 1a (Su(var)2-5) (Vermaak and Malik, 2009, Klattenhoff et al., 2009). While Su(var)2-5 mediates gene silencing and chromatin compaction, Rhino acts in an opposite manner: It enables the recruitment of several germline-specific proteins, which are required to engage the cellular gene expression machinery at piRNA clusters and enable the nuclear export of the emerging transcripts (ElMaghraby et al., 2019, Andersen et al., 2017, Mohn et al., 2014, Kneuss et al., 2019, Zhang et al., 2014b, Zhang et al., 2012a, Zhang et al., 2018, Hur et al., 2016, Chen et al., 2016). In *rhino* mutant flies, dual-strand piRNA clusters lose their transcriptional capacity and convert into canonical heterochromatin. Consequently, piRNA production collapses, transposons are de-repressed, and flies are sterile. Based on these data Rhino is the defining feature of dual-strand piRNA clusters, and its specific deposition is at the very center of steering piRNA populations to target ‘non-self’ (transposons) but not ‘self’ (genic loci).

Despite an increasing understanding of how Rhino orchestrates piRNA cluster expression, little is known about how cells control the specific deposition of Rhino onto chromatin. Current models involve a role of maternally inherited Piwi-piRNA complexes in the specification of piRNA cluster identity through Piwi-dependent deposition of H3K9 methylation, which is postulated to define dual-strand clusters at the chromatin level (Mohn et al., 2014, Le Thomas et al., 2014, Shpiz et al., 2014, Akkouche et al., 2017). In this manner, Rhino can be recruited adaptively to a changing transposon insertion profile. Indeed, Rhino’s N-terminal chromodomain displays specific affinity to H3K9me3, which is consistently present at piRNA clusters (Mohn et al., 2014, Le Thomas et al., 2014, Yu et al., 2015). However, a large fraction of the pericentromeric, H3K9me3 enriched heterochromatin is not or only weakly bound by Rhino. Since its discovery, a major open question has therefore been how Rhino’s genomic binding profile is defined.

Two observations indicate that, besides H3K9me3, cellular and genomic context impact Rhino’s binding pattern. First, while Rhino is expressed both in ovaries and testes, the identity of dual-strand piRNA clusters differs in the two gonads despite identical DNA sequence (Mohn et al., 2014, Klattenhoff et al., 2009, Chen et al., 2021). Moreover, the genomic Rhino profile is dynamic in testes where the X-chromosomal *AT-chX* cluster attracts most of the cellular Rhino pool in differentiating spermatocytes but not at earlier developmental stages (Chen et al., 2021). Second, while all stand-alone insertions of active transposons are silenced through piRNA-guided heterochromatin formation, only around 20% of the insertions of any given transposon are, by unknown mechanisms, bound by Rhino (Mohn et al., 2014, Shpiz et al., 2014, Radion et al., 2019, Akulenko et al., 2018). Based on these two observations, Rhino- domains must be defined through a combination of H3K9me3 with additional, unknown activities that bind chromatin via Piwi-independent mechanisms.

Here, we show that the zinc finger protein CG2678/Kipferl defines the majority of Rhino’s chromatin binding pattern in ovaries. The combinatorial readout of Kipferl’s sequence specific DNA binding together with Rhino’s affinity to the H3K9me3 mark underlies the selective recognition of heterochromatic loci as substrates for piRNA precursor transcription. Our findings also suggest that distinct chromatin loci might compete for the cellular Rhino pool, offering insight into why Rhino is a fast-evolving protein.

## RESULTS

### The H3K9me3 mark alone cannot explain the large diversity of genomic Rhino domains

As a foundation for studying how Rhino’s chromatin binding specificity is defined molecularly, and how variable it is in fly strains with different transposon insertion profiles, we determined Rhino’s chromatin occupancy genome-wide in two commonly used laboratory strains (*w^1118^* and *MTD-Gal4*) and in *iso1*, the strain underlying the *D. melanogaster* reference genome (Hoskins et al., 2015). Chromatin immunoprecipitation experiments followed by next-generation sequencing (ChIP-seq) confirmed that Rhino binds mostly to extended domains that vary greatly in size and often lack clearly defined boundaries (Mohn et al., 2014). We therefore divided the genome into nonoverlapping genomic 1-kb tiles and calculated the Rhino enrichment per tile in each strain. This revealed that a substantial fraction of the genome (e.g. 3.9 Mbp corresponding to ∼3% of the analyzable genome in the *w^1118^* strain) showed greater than 4-fold enrichment (p = 0.036, Z-score = 2.1) for Rhino over input.

To analyze Rhino’s chromatin binding profile, we divided the genome into pericentromeric heterochromatin and the generally euchromatic chromosome arms (based on H3K9me3 and Su(var)2-5 ChIP-seq data) (Fig. 1A; Table S1). The three well-described Rhino-dependent dual-strand piRNA clusters – *cluster 38C*, *42AB*, and *80F –* were not assigned to either category but were analyzed separately as reference loci. Rhino enrichment at major dual-strand clusters but also within the recombination-poor, pericentromeric heterochromatin was highly similar in all three fly strains (Fig. 1B left, middle; Fig. S1A). Many Rhino domains in pericentromeric heterochromatin (e.g. Fig. 1C panel 1) resembled the large piRNA clusters like *cluster 80F* (Fig. 1C panel 2): they exhibited strong H3K9me3 signal, gave rise to abundant piRNAs, and were enriched in diverse transposon sequences. Within chromosome arms, several strain-specific Rhino domains were apparent (Fig. 1B right). Based on manual inspection, these corresponded to genomic loci flanking stand-alone transposon insertions (e.g. a *flea* insertion found only in the *w^1118^* strain (Fig. 1C panel 3)). The majority of euchromatic Rhino domains, however, were shared among all three strains (Fig. 1B right; Fig. S1A). Based on the *iso1* data and the reference genome, many of these shared domains were not associated with transposon insertions. Upstream of the *headcase* locus, for example, Rhino was strongly enriched in all strains, although this locus lacks transposon sequences and does not produce abundant piRNAs (Fig. 1C panel 4). Nonetheless, these loci displayed dual-strand transcription based on RIP-seq data for Nxf3 (ElMaghraby et al., 2019), the piRNA precursor specific nuclear export factor, indicating that these loci were functional Rhino domains (Fig. 1C). The poor piRNA output from these transposon-devoid loci is therefore not due to non-functional Rhino, but most likely due to a lack of piRNA target sites for Aub/Ago3-mediated transcript cleavage in the emerging transcripts, which greatly stimulate piRNA biogenesis (Mohn et al., 2015, Han et al., 2015). Altogether, our data demonstrate that Rhino, in a largely strain-independent manner, binds to thousands of remarkably diverse genomic loci that include, but are not limited to sources of dual-strand piRNAs.

**Fig. 1:**
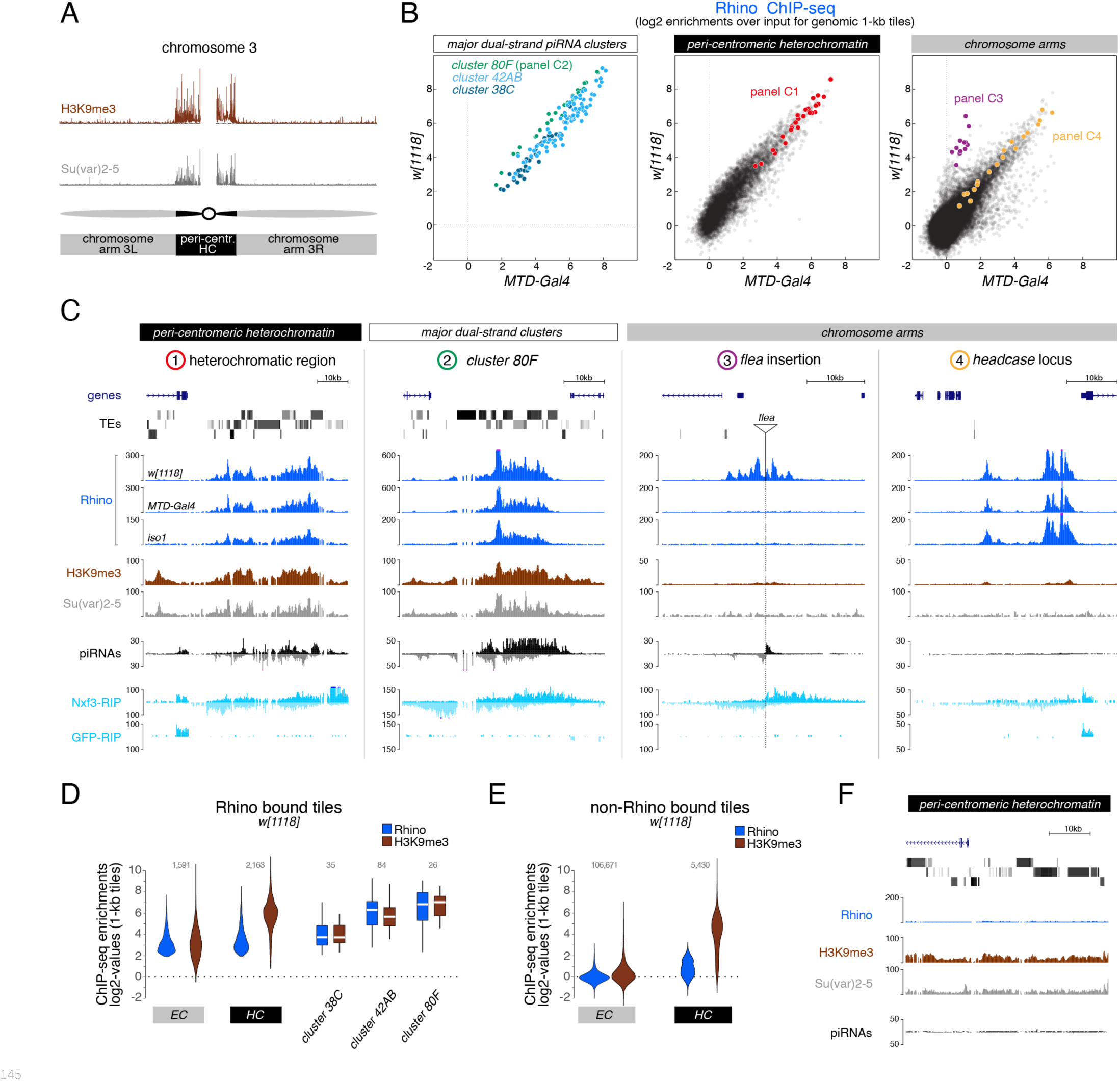
H3K9me3 is insufficient to define the large diversity of genomic Rhino domains. **(A)** ChIP-seq enrichment (genome unique reads; 1-kb tiles) of H3K9me3 and Su(var)2-5 along the assembled chromosome 3 sequence in *w[1118]* ovaries. Pericentromeric heterochromatin and euchromatic chromosome arms are indicated. **(B)** Scatter plot comparing average log2 ChIP-seq enrichments for Rhino in ovaries from *w[1118]* (n=2) versus MTD-Gal4 > *w*sh (n=3) strains (1-kb tiles separated into pericentromeric heterochromatin and chromosomal arms; piRNA clusters *38C*, *42AB*, and *80F* are shown separately; colored 1-kb tiles correspond to example loci in panel C). **(C)** UCSC genome browser tracks depicting the diversity of Rhino domains. F1: heterochromatic, transposon-rich locus; F2: piRNA *cluster 80F*; F3: strain-specific *flea* insertion (ElMaghraby et al., 2019) in *w[1118]*; F4: Rhino domain proximal to euchromatic *headcase* locus. Unless indicated otherwise, data are from *w[1118]* ovaries (ChIP and RIP signal are depicted as coverage per million reads, piRNA coverage normalized to miRNA reads). GFP-RIP-seq serves as control for non-specific mRNA binding. Genomic coordinates of individual panels are listed in Table S3. **(D, E)** Violin plots showing average log2 fold enrichment of Rhino ChIP-seq over input for 1-kb tiles (two replicate experiments) from *w[1118]* ovaries. Tiles were grouped into Rhino-bound and non-Rhino-bound based on a cutoff of 4-fold enrichment (corresponding to p = 0.036, Z-score = 2.1) of Rhino ChIP-seq signal over input in each replicate experiment. Rhino-dependent piRNA *clusters 38C*, *42AB*, and *80F* were analyzed separately (shown as box plots due to low number of tiles). **(F)** UCSC genome browser tracks depicting a pericentromeric heterochromatin locus marked by H3K9me3 and bound by Su(var)2-5, but not Rhino (ChIP-seq signal: coverage per million reads; piRNA coverage normalized to miRNA reads; all data obtained from *w[1118]* ovaries; genomic coordinates: Table S3).

Consistent with a reported role of H3K9 methylation in dual-strand piRNA cluster biology, all Rhino domains in both pericentromeric heterochromatin and within chromosome arms showed enrichment in H3K9me3 (Fig. 1D). However, H3K9me3 levels did not directly correlate with Rhino enrichments: First, enrichments of H3K9me3 were considerably higher for Rhino domains within pericentromeric heterochromatin compared to those in chromosome arms, despite comparable Rhino enrichments (Fig. 1D). Second, numerous loci in heterochromatin with high H3K9me3 signal were not or only poorly bound by Rhino (Fig. 1E, F). The recombinant Rhino chromodomain also binds to H3K9me2, yet with lower affinity (Mohn et al., 2014, Le Thomas et al., 2014, Yu et al., 2015). As for H3K9me3, however, a large fraction of H3K9me2 was found outside of Rhino-bound loci (Fig. S1B). In contrast, Su(var)2-5 enrichments correlated with H3K9me2/3 levels at almost all genomic loci (Fig. S1C compared to Fig. S1B). This was also true for a tagged Su(var)2-5 protein expressed exclusively in germline cells where it competes with Rhino for binding to H3K9me3. Thus, methylation of H3K9 alone, while being a hallmark of Rhino domains, cannot explain Rhino’s chromatin profile.

### The ZAD-zinc finger CG2678 interacts with Rhino and binds Rhino domains genome-wide

Sequence-specific DNA binding proteins can guide HP1 family proteins to chromatin. HP1b and HP1c, for example, depend on the zinc finger proteins Woc and Row for chromatin binding to thousands of sites that lack signal for Su(var)2-5 and even H3K9me2/3 (Fig. S1D) (Font-Burgada et al., 2008). Considering this, we hypothesized that undiscovered proteins with affinity to DNA motifs or to other histone marks collaborate with H3K9me3 to determine Rhino’s genomic target loci.

To identify Rhino-interacting proteins, we performed a yeast two hybrid screen using full length Rhino as bait and a cDNA library obtained from ovary mRNAs as prey (Fig. 2A). We identified 26 putative interactors among 175 sequenced colonies (Table S2), among them the known Rhino-interactor Deadlock (83 independent clones) (Mohn et al., 2014). With twelve independent clones, the uncharacterized protein CG2678 was the second most enriched screen hit. CG2678 further stood out among the other identified interactors because of its domain architecture: *CG2678* encodes a predicted DNA binder whose two annotated protein isoforms carry one or two arrays of C_2_H_2_ zinc fingers (ZnFs) (Fig. 2B). Based on an N-terminal ZnF associated domain (ZAD), CG2678 is a member of the large group of ZAD-ZnF proteins that have radiated extensively in insects (Chung et al., 2007, Chung et al., 2002). Only a handful of the >90 *D. melanogaster* ZAD-ZnF proteins have been studied, and these are involved in transcription (Bag et al., 2021, Harms et al., 2000), genome organization (Maksimenko et al., 2015, Sabirov et al., 2021), and heterochromatin biology (Kasinathan et al., 2020). As a predicted DNA-binding protein and a putative Rhino interactor, CG2678 was an intriguing candidate for a potential Rhino specificity factor.

**Fig. 2:**
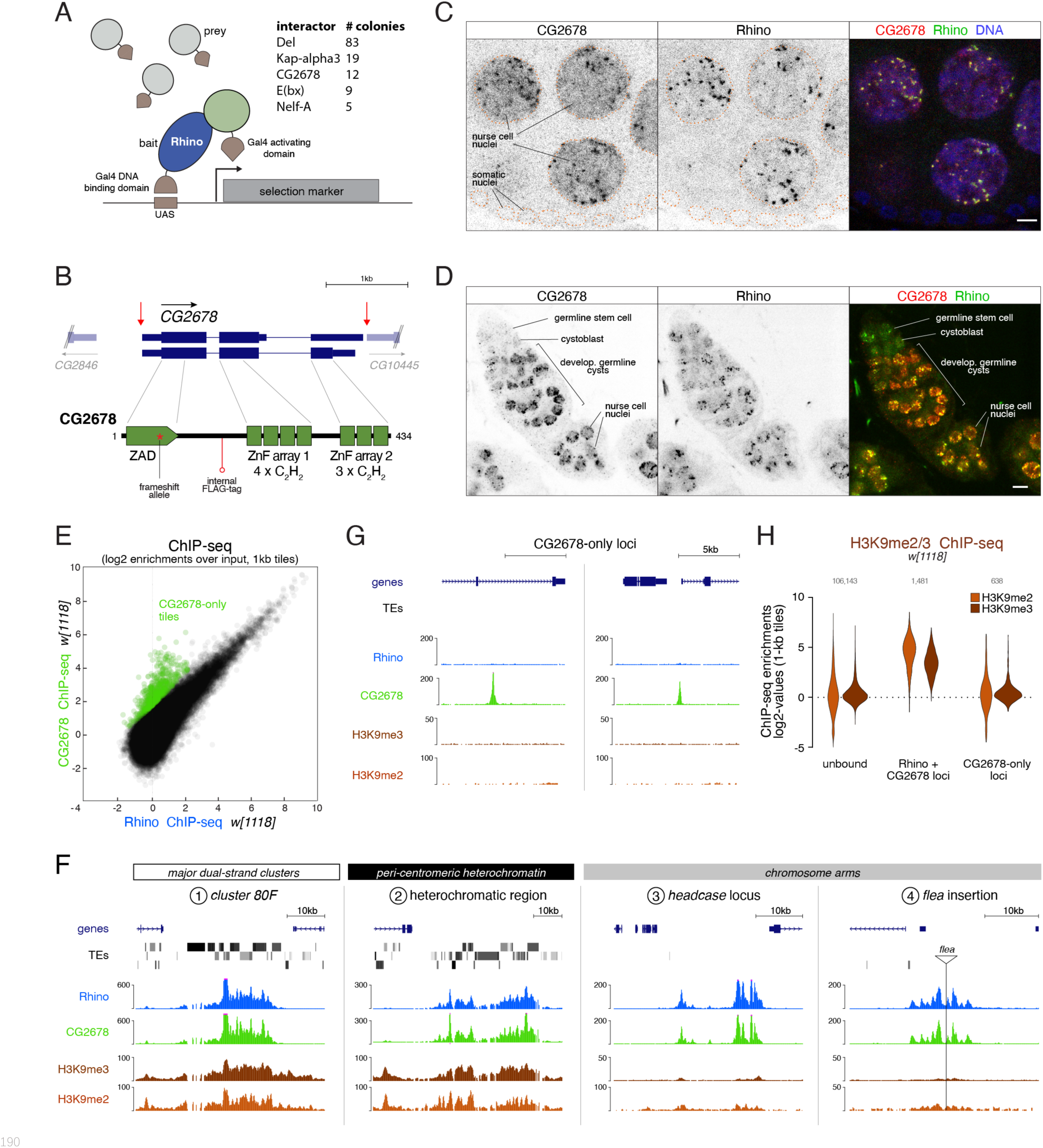
The ZAD-zinc finger CG2678 interacts with Rhino and binds Rhino domains genome-wide. **(A)** Cartoon illustrating yeast two hybrid screen setup for Rhino. Screen hits recovered >5 times are listed. **(B)** Genomic *CG2678* locus depicting the two annotated transcripts and CG2678 domain architecture (location of the frame shift mutation (red asterisk), internal 3xFLAG affinity tag (red circle), and cleavage sites for full locus deletion (red arrows) are indicated). **(C, D)** Confocal images of nurse cell nuclei (C) and germarium (D). Single channel and merged color images depict immunofluorescence signal for endogenous CG2678 (left, red) and Rhino (middle, green). Scale bar: 5 µm, dotted line: nuclear outline based on DAPI. **(E)** Scatter plot depicting correlation of log2-fold Rhino versus CG2678 ChIP-seq enrichment in *w[1118]* ovaries (average of 2 replicates each). CG2678-only tiles are highlighted in green and were defined by significantly higher enrichment for Rhino than CG2678 (n=2, Z-score = 3), plus a Rhino enrichment of max. 4-fold in two independent experimental replicates. **(F, G)** UCSC browser tracks illustrating CG2678 signal at diverse Rhino domains (F) and at CG2678-only peaks (G). ChIP-seq signal is shown as coverage per million sequenced reads for *w[1118]* ovaries (genomic coordinates of individual panels listed in Table S3). **(H)** Violin plots depicting log2-fold enrichment of H3K9me2 (orange) and H3K9me3 (brown) at 1-kb tiles bound by neither Rhino nor CG2678, both proteins, or CG2678 only. Classification into groups was performed based on binary cutoffs for Rhino (4-fold) and a linear fit for CG2678 co-occupancy in two independent replicate ChIP-seq experiments from *w[1118]* ovaries to extract CG2678-only tiles highlighted in (E).

Like for *rhino*, *CG2678* mRNA levels were detected primarily in ovaries (Flybase gene expression atlas, Fig. S2A) (Larkin et al., 2021). Immuno-fluorescence experiments revealed CG2678 expression specifically in ovarian germline cells, with no detectable signal above background in somatic cells which lack Rhino-defined piRNA clusters (Fig. 2C). In germline nurse cells, CG2678 was enriched in nuclei and accumulated in numerous discrete foci in a pattern that was almost indistinguishable from that of a Rhino staining (Fig. 2C). Expression of CG2678 in ovaries, however, was not uniform: Single cell RNA-seq data from adult ovaries indicated low mRNA levels in germline stem cells and increased levels in differentiating nurse cells (Rust et al., 2020). Consistent with this, CG2678 protein, while abundant in differentiating cysts and polyploid nurse cells, was barely detectable in germline stem cells and cystoblasts despite these cells expressing Rhino (Fig. 2D). CG2678 was further not detectable in testes (Fig. S2A, B). These data were intriguing given that the identity of Rhino-dependent dual-strand piRNA clusters differs, for unknown reasons, between males and females (Chen et al., 2021).

ChIP-seq experiments using anti-CG2678 antibody confirmed that the co-localization of CG2678 and Rhino in nuclear foci was due to both proteins binding the same chromatin sites, as CG2678 co-occupied Rhino domains genome-wide in all three wildtype strains (*w^1118^*, *MTD-Gal4*, and *iso1* strains) (Fig. 2E; Fig. S2C, D). Closer inspection revealed that CG2678 and Rhino bind to chromatin in a virtually indistinguishable pattern, often occupying broad chromatin domains. This was true at major piRNA clusters (e.g. *cluster 80F*), Rhino domains in pericentromeric heterochromatin, stand-alone transposon insertions in chromosome arms as well as loci like *headcase* that lack transposon sequences and piRNA output (Fig. 2F).

Notably, 647 genomic 1-kb tiles exhibited significantly higher enrichment for CG2678 than for Rhino (Z-score > 3) and were not enriched for Rhino (CG2678-only loci; 15.4% of all Kipferl-bound tiles) (Fig. 2E). At these loci, the CG2678 ChIP-seq signal formed narrow peaks (e.g. Fig. 2G). 98.7% of the CG2678-only loci resided within chromosome arms, suggesting that lack of H3K9 methylation at these sites might prevent Rhino binding. Indeed, almost all CG2678-only loci were devoid of H3K9 methylation (Fig. 2G, H). Overall, our findings revealed a molecular relationship between Rhino and the potentially sequence-specific DNA binder CG2678.

### Rhino’s chromatin occupancy changes dramatically in *CG2678/kipferl* mutants

To investigate the role of CG2678 in Rhino biology, we generated mutant fly lines carrying frameshift alleles or a complete deletion of the *CG2678* locus (Fig. 2B). *CG2678* null mutant flies were viable and, in Western blot analysis, did not express detectable CG2678 protein (Fig. 3A). *CG2678* mutant females showed normal egg-laying rates but strong fertility defects, which were elevated in older females (Fig. 3B; Fig. S3A). Insertion of the *CG2678* genomic sequence with an internal FLAG-tag (Fig. 2B) at the deleted locus restored fertility to wildtype levels (Fig. 3B).

**Fig. 3:**
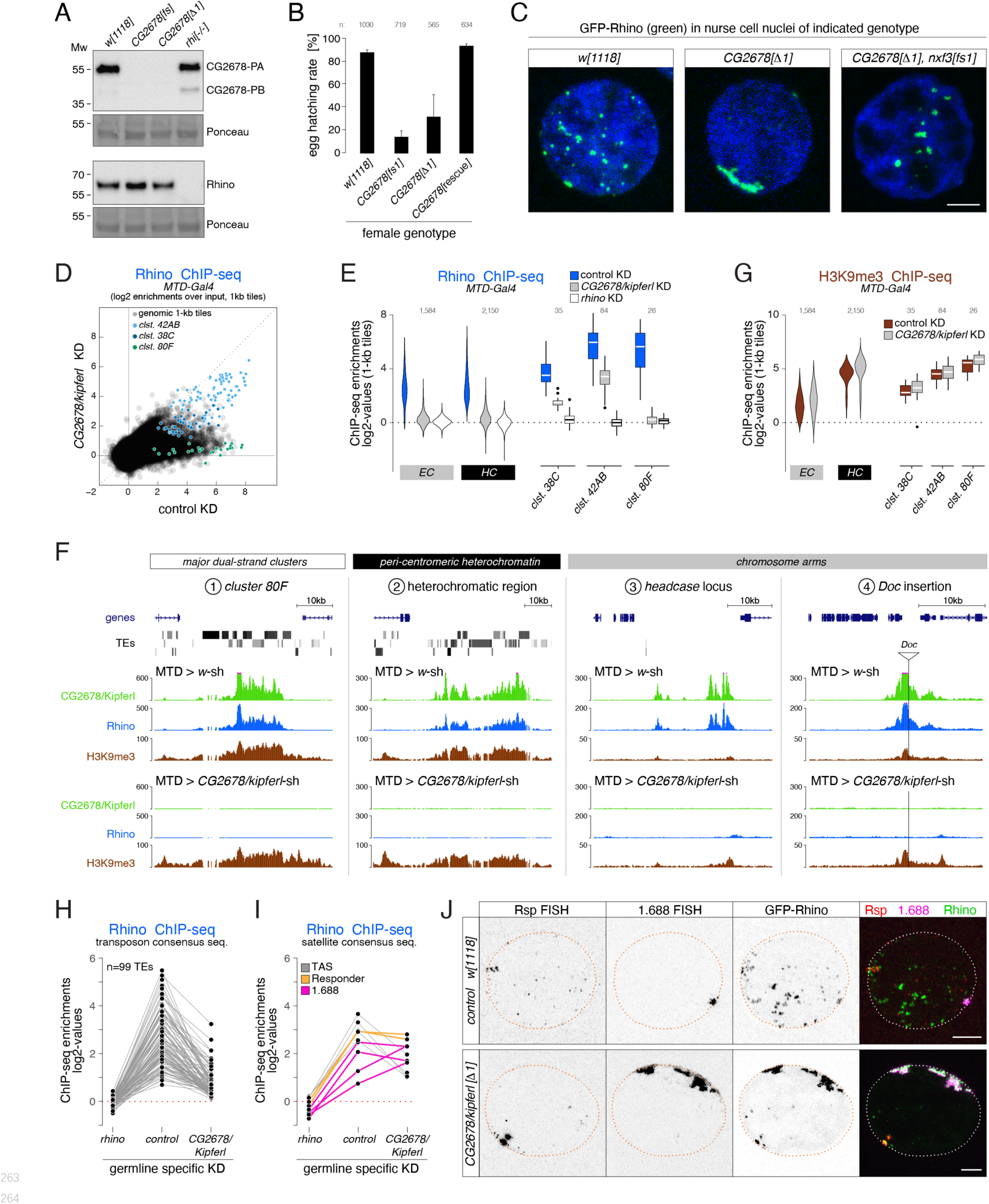
Rhino’s chromatin occupancy changes dramatically in *CG2678/kipferl* mutants. **(A)** Western blot analysis verifying *CG2678* frame shift (fs1) and locus deletion (Δ1) alleles using a monoclonal antibody against CG2678 (top; CG2678-PB is a minor protein isoform) and depicting Rhino levels in the absence of CG2678 (bottom). Ponceau staining: loading control. **(B)** Bar graph indicating average embryo hatching rate of eggs laid by *w[1118]* control females in comparison to females carrying a *CG2678* frame shift (fs1), locus deletion (Δ1), or *CG2678* null mutants with a tagged rescue construct, respectively (average of four time points collected over a 2 day period; see also Fig.S3A; total number of eggs laid indicated). **(C)** Confocal images illustrating localization of GFP-Rhino in nurse cell nuclei of *w[1118]*, *CG2678* locus deletion (Δ1), and *CG2678,nxf3* double mutant females (scale bar: 5 µm). **(D)** Scatter plot of genomic 1-kb tiles contrasting average log2-fold Rhino ChIP-seq enrichment in ovaries with MTD-Gal4 driven *CG2678/kipferl* knock down versus control ovaries (two replicate experiments each). **(E)** Violin plots showing average log2-fold Rhino ChIP-seq enrichment in indicated germline knockdown ovaries on Rhino-bound 1-kb tiles (defined in Fig. 1D) in heterochromatin (HC) and chromosome arms (EC). piRNA clusters *38C*, *42AB*, and *80F* are depicted separately. **(F)** UCSC browser tracks (ChIP-seq) depicting diverse Rhino domains in control and *CG2678/kipferl* germline knockdown ovaries (signal shown as coverage per million sequenced reads; genomic coordinates of individual panels listed in Table S3). **(G)** Violin plots showing average log2-fold H3K9me3 ChIP-seq enrichment in indicated genetic backgrounds for Rhino-bound 1-kb tiles (defined in Fig. 1D) in heterochromatin (HC) and along chromosome arms (EC). piRNA clusters *38C*, *42AB*, and *80F* are depicted separately. **(H, I)** Jitter plots depicting the log2-fold Rhino ChIP-seq enrichments on transposon (H) and Satellite (I) consensus sequences in indicated genetic backgrounds. **(J)** Confocal images showing *Rsp* and *1.688* Satellite RNA FISH signal and GFP-Rhino in nurse cells of *w[1118]* or *CG2678/kipferl* mutant flies (scale bar: 5 µm).

Loss of CG2678 had no impact on Rhino levels (Fig. 3A) but resulted in pronounced changes in Rhino localization: In wildtype nurse cells, Rhino accumulated in many distinct foci throughout the nucleus. In *CG2678* mutants, almost all Rhino signal gathered in a few large, often continuous structures at the nuclear envelope (Fig. 3C). Depletion of *CG2678* mRNA by germline-specific RNAi caused a similar phenotype (Fig. S3B, C), and expression of a tagged wildtype CG2678 protein in *CG2678* mutants restored wildtype Rhino localization to small, dispersed foci (Fig. S3D). The prominent Rhino accumulations in *CG2678* mutants were enriched in H3K9me3 and the Rhino co-factors Deadlock and Nxf3, indicating that they were genuine chromosomal Rhino domains (Fig. S3E).

Rhino domains tend to localize to the nuclear envelope also in wildtype ovaries, resulting in a putative piRNA precursor export and biogenesis compartment continuous with cytoplasmic nuage (Zhang et al., 2012a). We hypothesized that the pulling force provided by the piRNA precursor export pathway centered on Nxf3 drags Rhino domains to nuclear pore complexes (ElMaghraby et al., 2019). Indeed, in *CG2678*,*nxf3* double mutant flies, Rhino still accumulated in large foci but these were no longer confined to the nuclear envelope (Fig. 3C). We termed *CG2678 ‘kipferl’* because of the prominent enrichments of Rhino in crescent-shaped structures that reminded us of a famous Austrian pastry.

To explore whether Rhino’s altered nuclear localization was linked to changes in its chromatin occupancy, we performed Rhino ChIP-seq experiments in *kipferl*-depleted ovaries. In the absence of Kipferl, Rhino’s chromatin association was severely reduced at most genomic sites (Fig. 3D, E). These included the major piRNA cluster *80F*, nearly all Rhino domains in pericentromeric heterochromatin, as well as stand-alone transposon insertions and euchromatic Rhino domains devoid of transposon insertions (Fig. 3E, F). A similar phenotype was apparent in ovaries from flies carrying *kipferl* null mutant alleles (Fig. S3F). Importantly, Rhino loss at these various genomic loci was not connected to reduced H3K9 methylation levels (Fig. 3G; Fig. S3G).

Only 4.6% of all 1-kb tiles exhibiting a more than 4-fold Rhino enrichment (n=2) in control flies remained Rhino-bound in *kipferl*-depleted ovaries. Among these were mainly tiles of piRNA *cluster 42AB* and, to a lesser extent, *cluster 38C* (Fig. 3D). Fluorescence *in situ* hybridization (FISH) experiments confirmed that *cluster 42AB* and *38C* are transcribed in *kipferl* mutants (Fig. S3H). However, their respective nuclear RNA-FISH signal did not co-localize with the strong, elongated Rhino accumulations at the nuclear envelope (Fig. S3H). We therefore hypothesized that the prominent Rhino foci in *kipferl* mutants correspond to repetitive loci not assembled in the reference genome or not identifiable using genome-unique reads. Based on an analysis of all ChIP-seq reads, the average Rhino enrichment on transposon consensus sequences varied between two and 30-fold in wildtype ovaries, was non-detectable in ovaries depleted for Rhino, and was strongly reduced in the absence of Kipferl (Fig. 3H). In contrast, several Satellite sequences, foremost the *Responder* and *1.688* family Satellites that give rise to Rhino-dependent piRNAs in wildtype ovaries and testes (Chen et al., 2021, Wei et al., 2021), maintained Rhino occupancy or accumulated even more Rhino than in wildtype ovaries (Fig. 3I).

Consistent with elevated Rhino occupancy at both Satellites, we observed a corresponding increase in nascent transcripts as indicated by the enhanced RNA FISH-signal in nurse cell nuclei of *kipferl* mutant ovaries compared to wildtype controls (Fig. 3J). The FISH signal for *Rsp* and *1.688* transcripts in *kipferl* mutants overlapped precisely with the prominent Rhino accumulations at the nuclear envelope. *Rsp* and *1.688* Satellites form large repetitive tandem arrays in pericentromeric heterochromatin (Wu et al., 1988, Abad et al., 2000). The enormous size of these Satellite loci, particularly the *359bp* array (*1.688* family) on the X-chromosome which extends over several megabasepairs, likely explains why Rhino foci in *kipferl* mutants are so prominent (Fig. 3J). In summary, Rhino’s re-distribution from hundreds of genomic loci to the large, pericentromeric *Rsp* and *1.688* satellite arrays, which accumulate at the nuclear periphery in an Nxf3-dependent manner (Fig. S3I), explain the name-giving Rhino localization phenotype in *kipferl* mutants.

### *kipferl* mutant ovaries display piRNA losses and transposon de-repression

To investigate whether Rhino’s altered chromatin occupancy in *kipferl* mutants affects piRNA production, we compared Argonaute-bound small RNA populations from *kipferl*-depleted ovaries to those from *rhino*-depleted and control ovaries. Total piRNA levels, normalized to miRNA reads, were reduced 4.5-fold in *rhino*-depleted but only 1.5-fold in *kipferl*-depleted ovaries (Fig. S4A). We grouped genomic piRNA sources into somatic source loci (e.g. *flamenco*), Rhino-independent germline source loci (e.g. uni-strand *cluster 20A*) and Rhino-dependent germline source loci (e.g. dual-strand *clusters 38C*, *42AB*, *80F*) (Fig. S4B) (Mohn et al., 2014). *kipferl*-depleted ovaries exhibited reduced piRNA levels only from Rhino-dependent germline source loci (Fig. 4A). A prominent example was *cluster 80F*, where piRNA production collapsed in the absence of Kipferl to the same extent as seen in *rhino*-depleted ovaries (Fig. 4A, B). At many other sites, piRNA losses were less severe compared to *rhino*-depleted ovaries. Among them were piRNA *clusters 38C* and *42AB*, consistent with residual Rhino binding to both clusters in *kipferl*-depleted ovaries (Fig. 4A, B; Fig. 3D, E).

**Fig. 4:**
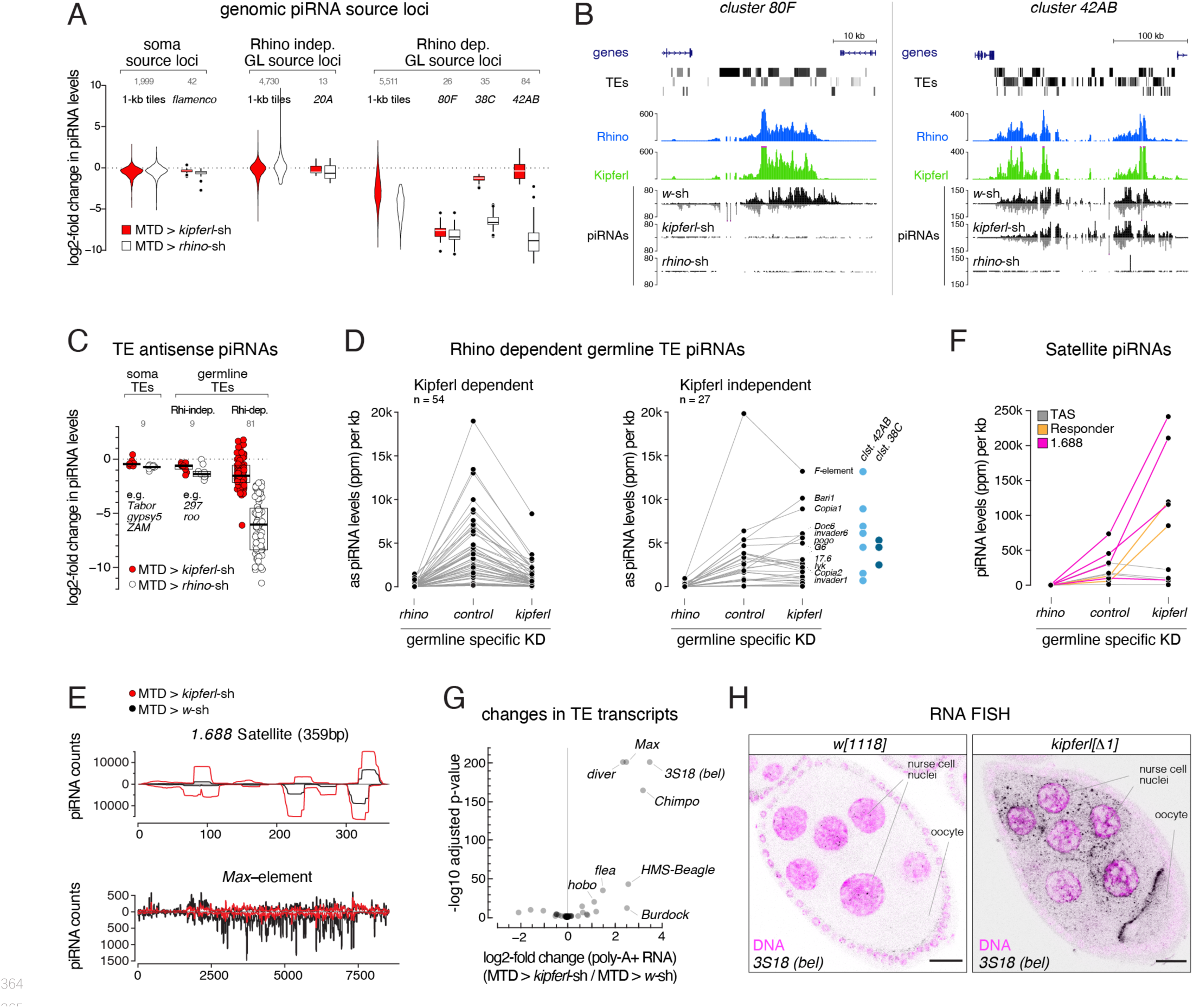
*kipferl* mutant ovaries display piRNA losses and transposon de-repression. **(A)** Violin plots showing log2-fold changes in levels of uniquely mapping piRNAs on 1-kb windows relative to control upon MTD-Gal4 mediated knock down of *rhino* or *kipferl* (1-kb tiles were categorized into somatic source loci, Rhino-independent germline source loci and Rhino-dependent germline source loci according to Fig. S4B). **(B)** UCSC genome browser tracks displaying piRNA levels at cluster *80F* and *42AB* in control, *kipferl*, and *rhino* knock down ovaries (ChIP-seq signal from MTD-Gal4 control ovaries is depicted as coverage per million reads, piRNA coverage was normalized to miRNA reads; genomic coordinates of individual panels listed in Table S3). **(C)** Jitter plots depicting log2-fold changes for piRNA levels mapping antisense to transposon consensus sequences in indicated MTD-Gal4 mediated knock downs compared to control (transposons classified analogous to panel A). **(D, E)** Jitter plots showing piRNA levels (per kb sequence) in indicated genotypes mapping to transposons (antisense only) giving rise to Rhino-dependent piRNAs (D) or to Satellite repeats (E). Blue dots in panel D indicate insertions of the respective transposon in piRNA clusters *38C* or *42AB*. **(F)** piRNA profiles across the consensus sequences for a representative transposon (*Max*) and the 359bp *1.688* Satellite. piRNA counts (normalized to miRNAs) are displayed for indicated genotypes. **(G)** Volcano plot depicting the log2-fold changes in poly-adenylated transposon transcripts in *kipferl* depleted versus control ovaries (n=3). **(H)** Confocal images showing RNA FISH signal for *3S18* transcripts in *w[1118]* and *kipferl* null mutant ovaries (scale bar: 20 µm).

The selective impact of *kipferl* depletion on germline-specific, Rhino-dependent piRNA source loci was also reflected when piRNAs mapping in antisense orientation to transposon consensus sequences were analyzed (Fig. 4C). piRNAs targeting soma-controlled transposons (e.g. *Tabor*, *gypsy5*, *ZAM*; mostly originating from *flamenco*) were not affected in *rhino*- or *kipferl*-depleted ovaries. Among the transposons with dominating germline piRNA populations, nine are targeted by Rhino- and Kipferl- independent piRNAs originating from Rhino-independent source loci like *cluster 20A* (e.g. *297*, *roo*). All other transposons were targeted by Rhino-dependent piRNAs. For most of these elements, piRNA pools were reduced in *kipferl*-depleted ovaries (Fig. 4D left, E). The few transposons that retained piRNAs or even displayed increased piRNA levels had insertions in piRNA *clusters 38C* or *42AB*, where Rhino binding was largely Kipferl-independent (Fig. 4D right).

We finally analyzed the *TAS*, *Rsp*, and *1.688* Satellite repeats, which give rise to Rhino-dependent piRNAs (Fig. 4F) (Chen et al., 2021, Wei et al., 2021). While piRNA levels from sub-telomeric *TAS* Satellites remained unchanged, those derived from the *Rsp* and *1.688* Satellites increased strongly in *kipferl*-depleted ovaries compared to control, consistent with increased Rhino occupancy at these loci (Fig. 4E, F; Fig. 3I). As a result, Satellite piRNAs which accounted for only 4% of all piRNAs in wildtype ovaries, accounted for more than one in five piRNAs (21%) in *kipferl*-depleted ovaries. Thus, in the absence of Kipferl, redistribution of Rhino from its natural binding sites to pericentromeric Satellite repeats results in a substantial loss of transposon targeting piRNAs and increased Satellite piRNAs.

In line with reduced levels of piRNAs antisense to transposon sequences, the levels of poly-adenylated transcripts for a number of transposable elements increased significantly in *kipferl*-depleted ovaries (Fig. 4G). RNA-FISH experiments confirmed de-repression of these elements in *kipferl* null mutant ovaries (Fig.4H; Fig. S4C). We note that transposon de-repression in ovaries lacking Kipferl was less severe compared to ovaries lacking Rhino (Mohn et al., 2014). This might be due to the milder loss of piRNAs in *kipferl* mutants compared to *rhino* mutants, or due to Kipferl being expressed only upon germline cell differentiation.

Our combined genetic data suggested that Kipferl acts specifically in the piRNA pathway. Consistent with this, loss of Kipferl had hardly any effect on overall gene expression, similar to the piRNA-specific factor Rhino (Fig. S4D). By defining most of Rhino’s genomic binding sites and by preventing aberrant Rhino accumulations at Satellite repeats, Kipferl enables nurse cells to mount a robust and diverse piRNA defense that is essential for the tight repression of transposable elements. Based on these findings, we set out to understand the molecular connections between Kipferl, Rhino, and DNA/chromatin.

### Kipferl nucleates Rhino domains within H3K9me2/3 loci and binds guanosine-rich DNA motifs

Kipferl was enriched at all Rhino domains but also at hundreds of loci that lacked Rhino occupancy (Fig. 2E, G). At Rhino domains, Kipferl and Rhino enrichments were indistinguishable and covered broad chromatin domains, while at Kipferl-only sites, Kipferl was enriched in narrow peaks (Fig. 5A). We hypothesized that Kipferl is a direct, sequence-specific DNA binder that functions upstream of Rhino and recruits and/or stabilizes Rhino on chromatin at its binding sites if they are located within an H3K9me2/3 domain. To test this, we investigated Kipferl’s chromatin binding capacity in the absence of Rhino. In *rhino* mutant ovaries, Kipferl was distributed throughout nurse cell nuclei rather than being enriched in discrete nuclear foci as in wildtype ovaries (Fig. 5B). ChIP-seq experiments revealed that Kipferl nevertheless bound to thousands of chromatin sites in *rhino* mutants where it was enriched in narrow peaks akin to Kipferl-only loci in wildtype ovaries (Fig. 5C). No ChIP-seq signal was observed in ovaries from *kipferl* null mutants (Fig. S5A).

**Fig. 5:**
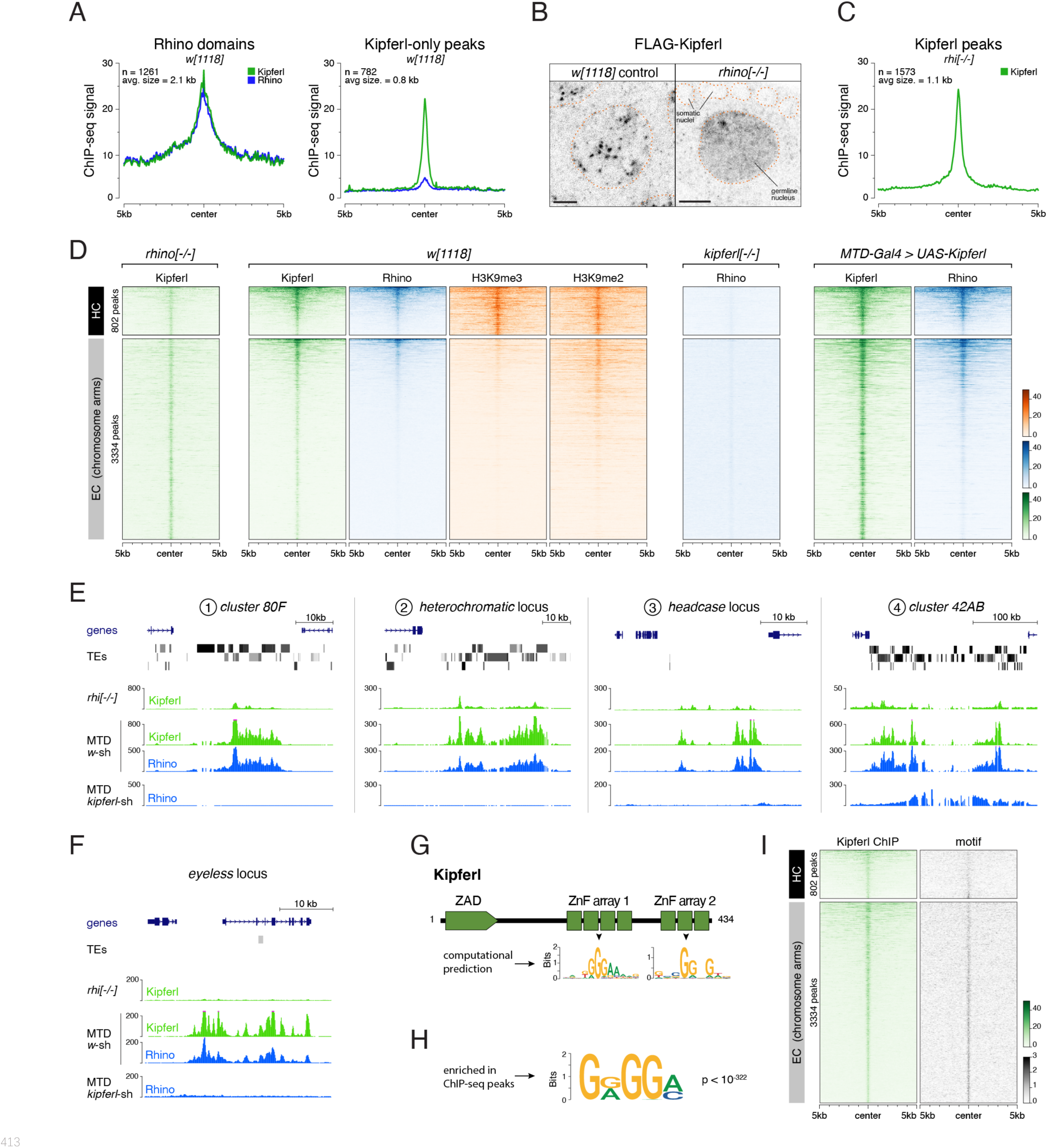
Kipferl binds guanosine-rich DNA motifs and nucleates Rhino domains within H3K9me2/3 loci. **(A)** Meta profiles showing the indicated ChIP-seq signal on Kipferl peaks co-occupied by Rhino (left) or bound by Kipferl only (right) in *w[1118]* ovaries (ChIP-seq signal is depicted as coverage per million reads). **(B)** Confocal images of nurse cell nuclei expressing FLAG-tagged Kipferl in indicated genetic backgrounds (scale bar: 5 µm; nuclear outlines based on DAPI as orange dotted line). **(C)** Meta profile showing Kipferl peaks determined in *rhino* mutant ovaries using the same peak calling parameters as in panel A. **(D)** Heat maps depicting indicated ChIP-seq signal in indicated genotypes from representative replicate experiments centered on narrow Kipferl peaks detected in two independent ChIP-seq experiments of *rhino* mutant ovaries (data sorted by Rhino signal in *w[1118]*). **(E, F)** UCSC genome browser tracks depicting Kipferl and Rhino ChIP-seq signals in indicated genotypes for diverse Rhino domains with (F) or without (G) pronounced Kipferl nucleation sites (ChIP-seq signal depicted as coverage per million reads; piRNA coverage normalized to miRNA reads; genomic coordinates of individual panels listed in Table S3). **(G)** Schematic representation of Kipferl protein domain architecture and binding motives predicted for the two ZnF arrays (Princeton Cys2His2 PWM predictor). **(H)** Position weight matrix of Kipferl consensus motif determined by HOMER from top 3,000 narrow Kipferl peaks found in two independent ChIP-seq replicates from *rhino* mutant ovaries. The GRGGN motif was found at least once in 83.4 % of peaks versus 50.3% in control sequences (p < 10^-322^). **(I)** Heat map depicting Rhino-independent Kipferl ChIP-seq signal and GRGGN motif enrichment centered on narrow Kipferl peaks analogous to panel E, sorted by Kipferl signal in *rhi[-/-]* (motif count: # of motifs per non-overlapping genomic 100 bp window).

Kipferl binding sites in *rhino* mutants were found both in pericentromeric heterochromatin (802 peaks) and within chromosomal arms (3334 peaks), and often coincided with regions bound by Kipferl and Rhino in wildtype ovaries, demonstrating that Kipferl binds the same sites in both genotypes, yet in different patterns (Fig. 5D). Kipferl’s chromatin enrichment expanded from the narrow peaks seen in *rhino* mutant ovaries to broad domains in wildtype ovaries, specifically at peaks where Rhino was present. Kipferl peaks that did not show Rhino binding, on the other hand, did not widen into extended domains and often decreased in intensity. This reflected our previous observations of narrow Kipferl- only sites and broad domains co-occupied by Kipferl and Rhino. Importantly, Rhino recruitment to Kipferl peaks correlated with H3K9me2/3 levels: Rhino occupied the majority of Kipferl peaks in pericentromeric heterochromatin where H3K9me2/3 is abundant. Within chromosomal arms, Rhino accumulated preferentially at those Kipferl sites that were within a local H3K9me2/3 domain. These findings supported a model in which Kipferl binds DNA independently of Rhino, Kipferl binding sites act as Rhino nucleation sites when inside local heterochromatin, and Kipferl and Rhino cooperate to spread from nucleation sites into flanking heterochromatic regions, resulting in extended Rhino/Kipferl domains. Consistent with this, Rhino occupancy at and around Kipferl binding sites was dependent on Kipferl and overexpression of Kipferl in germline cells resulted in increased Kipferl binding at all sites, leading to the strengthening of Rhino/Kipferl domains within H3K9me2/3 domains, and the formation of additional small Rhino domains at pre-existing Kipferl sites with low, but detectable H3K9me2/3 levels (Fig. 5D).

Kipferl’s intrinsic chromatin binding profile in *rhino* mutants was strongly predictive of Rhino’s chromatin occupancy, often mirroring the non-uniform enrichment of Rhino in wildtype ovaries, only at lower levels (Fig. 5E). At other loci (e.g. the *eyeless* gene), the extended Rhino domain in wildtype flies encompassed only rather weak putative Kipferl nucleation sites (Fig. 5F). Considering that Rhino was dependent on Kipferl also at these loci, we speculate that here, both proteins, supported by local H3K9- methylation, are required for formation of a stable Rhino/Kipferl domain. Finally, Kipferl influenced Rhino’s chromatin profile even at the largely Kipferl-independent piRNA *cluster 42AB*: Kipferl-independent Rhino-binding in the highly repetitive center of *42AB* was supported by Kipferl peaks towards the periphery, resulting in an extended Rhino domain in wildtype ovaries (Fig. 5E). Based on these data, we conclude that Kipferl is a major specificity factor for Rhino in ovaries, and that Kipferl cooperates with Rhino to form extended Rhino domains from defined nucleation sites. Our data further indicate that Kipferl is not the only Rhino specificity factor but demonstrate that it is also required for the stabilization of Rhino domains nucleated by alternative means.

According to the nucleation-site model, Kipferl is expected to bind to specific DNA motifs. Consistent with this, its two ZnF arrays are predicted to bind DNA with a specificity for guanosine-rich motifs (Fig. 5G) (Persikov and Singh, 2014). We determined sequence motifs that were enriched in Kipferl ChIP-seq peaks identified in *rhino* mutant ovaries. The top enriched motif (found in >80% of all peaks; p<10^-322^) closely matched the *in silico* predictions for Kipferl’s ZnF arrays (Fig. 5H). This GRGGN motif was locally enriched at experimentally determined Kipferl peaks, regardless of whether the peak was located in pericentromeric heterochromatin or chromosomal arms, or whether it was within a Rhino domain or constituted a Kipferl-only site (Fig. 5I). Additional motifs detected at lower frequency and confidence often displayed variations of the same guanosine-rich motif (Fig. S5B). In support of Kipferl’s specificity for a GRGGN sequence, stable overexpression of FLAG-tagged Kipferl in cultured ovarian somatic stem cells (OSCs), which do not express Kipferl or Rhino, resulted in Kipferl binding at thousands of defined sites that were enriched in the GRGGN sequence motif (Fig. S5C). Taken together, our data support a direct and specific Kipferl-DNA interaction at the basis of Rhino recruitment to chromatin.

### Structure-function analysis of Kipferl’s DNA and Rhino binding activities

The yeast two hybrid results (Fig. 2A) and ChIP-seq analyses (Fig. 5) suggested that Kipferl binds directly to both, Rhino and DNA. To examine how these two molecular activities are encoded within the Kipferl protein, we created flies that expressed truncated FLAG-tagged variants instead of endogenous Kipferl, lacking either one of the ZnF arrays (Kipferl^Δ-1st-array^, Kipferl^Δ-2nd-array^) or the N-terminal ZAD (Kipferl^ý-ZAD^) (Fig. 6A). Kipferl^Δ1st-array^ showed strongly reduced chromatin binding capabilities (Fig. 6B). Deletion of the second ZnF array (Kipferl^Δ2nd-array^) instead had only mild impacts. The N-terminal ZAD, which is characteristic for the >90 ZAD-ZnF family members in *D. melanogaster*, promotes anti- parallel homodimerization but has not been directly linked to DNA binding (Jauch et al., 2003, Bonchuk et al., 2021). Yeast two hybrid experiments confirmed the dimerization capability of Kipferl’s ZAD (Fig. S6A). Deletion of the ZAD, and therefore abrogation of Kipferl dimerization, resulted in a global loss of Kipferl ChIP-seq signal (Fig. 6B), consistent with studies about other ZAD-ZnF proteins (Maksimenko et al., 2020). Together, these findings indicated that Kipferl binds DNA as a dimer, primarily via its first ZnF array.

**Fig. 6:**
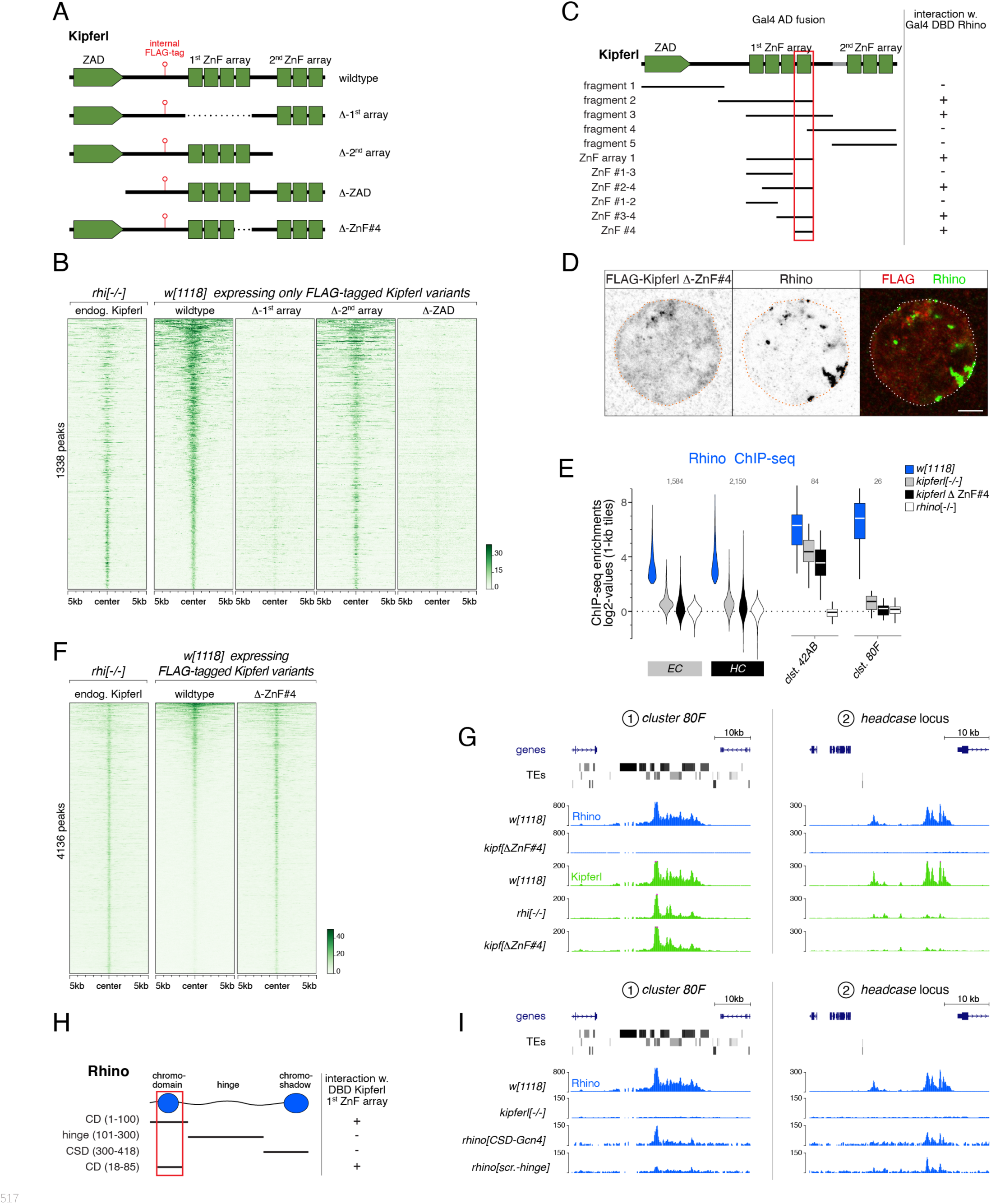
Structure-function analysis of Kipferl’s DNA and Rhino binding activities. **(A)** Schematic representation of Kipferl rescue constructs harboring the wildtype protein sequence or indicated deletions, as well as an internal 3xFLAG tag. Rescue constructs were introduced into the endogenous *kipferl* locus via RMCE. **(B)** Heat map displaying Kipferl variant ChIP-seq signal centered on peaks bound by Kipferl in *rhino* mutants (data sorted by the ChIP signal detected for the wildtype Kipferl rescue construct; shown are only peaks that are Kipferl-bound in ovaries expressing wildtype tagged Kipferl). **(C)** Schematic overview of yeast two hybrid experiments determining the minimal Rhino-interacting fragment in the Kipferl protein. Positive interactions between Kipferl fragments fused to the Gal4 activating domain (AD) and Rhino fused to the Gal4 DNA binding domain (DBD) are indicated. The minimal Kipferl fragment required for the Rhino interaction is highlighted in red (the grey bar indicates a 25 amino acid exon not contained in the reference genome which we identified in a subset of lab strains and included here for completeness). **(D)** Confocal images of a representative nurse cell nucleus depicting localization of Kipferl lacking ZnF#4 and Rhino in flies expressing only Kipferl-ΔZnF#4 (scale bar: 5 µm). **(E)** Violin plots showing the average log2-fold enrichment of Rhino ChIP-seq signal over input for Rhino-bound 1-kb tiles (classified in Fig. 1D) in the indicated genotypes. **(F)** Heat map showing indicated ChIP-seq signal, centered on narrow Kipferl peaks detected in two independent ChIP-seq experiments of *rhino* mutant ovaries (data sorted by ChIP-seq signal detected for wildtype Kipferl rescue). **(G)** USCS genome browser tracks showing ChIP-seq signal (coverage per million sequenced reads) for indicated proteins and genotypes at piRNA cluster *80F* (left) and the *headcase* locus (right). Genomic coordinates of individual panels listed in Table S3. **(H)** Schematic overview of yeast two hybrid experiments determining the minimal Kipferl-interacting fragment in Rhino. Positive interactions between Rhino fragments fused to the Gal4 activating domain (AD) and the first ZnF array of Kipferl fused to the Gal4 DNA binding domain (DBD) are indicated. The minimal Rhino fragment required for the Kipferl interaction is highlighted in red. **(I)** USCS genome browser tracks showing ChIP-seq signal (coverage per million sequenced reads) for indicated Rhino variants at piRNA cluster *80F* (left) or the *headcase* locus (right). Genomic coordinates of individual panels listed in Table S3.

To dissect the interaction of Kipferl with Rhino, we first examined the characteristic colocalization of Kipferl and Rhino in nurse cell nuclei. This revealed that both DNA binding defective variants, Kipferl^Δ1st-array^ and Kipferl^ΔZAD^, failed to colocalize with Rhino (Fig. S6B). In both genotypes, Rhino accumulated in prominent domains at the nuclear envelope, as seen in *kipferl* null mutant ovaries. Artificial dimerization of the Kipferl^ΔZAD^ variant via the heterologous dimerization domain from the yeast Gcn4 transcription factor or the ZAD of Ouija board, a ZAD-ZnF protein not expressed in ovaries, restored co-localization of Kipferl with Rhino (Fig.S6B, Kipferl^GCN4^ and Kipferl^ouib^). Thus, neither the ZAD, nor the second ZnF array are critical for binding to Rhino, suggesting that Kipferl’s 1^st^ ZnF array, besides its central role in DNA binding, might also enable Rhino binding.

Yeast two hybrid experiments, probing full length Rhino against Kipferl fragments, confirmed that Rhino interacts with Kipferl’s first ZnF array, with no additional interaction interfaces being identified (Fig. 6C, S6C). To disentangle the DNA-binding and Rhino-binding activities of the first ZnF array, we further narrowed down the interaction between Rhino and Kipferl, revealing a critical role of the 4^th^ ZnF (Kipferl^ýZnF#4^, Fig. 6C; Fig. S6C, D). We tested this putative split-of-function mutant *in vivo*. In flies expressing Kipferl^ýZnF#4^ instead of the endogenous protein, Rhino did not co-localize with Kipferl, and displayed the characteristic *kipferl* mutant phenotype (Fig. 6D; Fig. S6B). Moreover, Rhino lost its chromatin occupancy in *kipferl ^ΔZnF#4^* flies in a pattern also seen in *kipferl* null mutants (Fig. 6E). This included a complete loss of Rhino at Kipferl-dependent loci like *cluster 80F* or the *headcase* locus, but also remaining Rhino signal at Kipferl-independent loci like *cluster 42AB*, where Rhino’s ChIP-seq signal in *kipferl ^ΔZnF#4^* or *kipferl* null mutant flies was comparable (Fig. 6G; Fig. S6E). Kipferl^ýZnF#4^ was diffusely localized in nurse cell nuclei akin to wildtype Kipferl in *rhino* mutants, and *kipferl^ýZnF#4^* females were sub-fertile (Fig. 6D; Fig. S6F). Strikingly, Kipferl^ýZnF#4^ retained full DNA binding ability as it was enriched on chromatin in a pattern closely mirroring that of wildtype Kipferl in a *rhino* mutant (Fig. 6F). The Rhino- and DNA-binding activities of Kipferl can therefore be uncoupled: ZnFs 1-3, supplemented by the second ZnF array, allow for sequence specific DNA binding, while ZnF 4 interacts with Rhino.

The finding that the 4^th^ C_2_H_2_ ZnF fold in Kipferl is sufficient (Fig. 6C) and required (Fig. 6D, E) for Rhino binding was intriguing. Canonical HP1 interactors typically bind the chromoshadow domain dimer of HP1 proteins via PxVxL motif-containing peptides (Thiru et al., 2004). No such motif was found in Kipferl’s 4^th^ ZnF. We inverted the yeast two hybrid assay to determine which region of Rhino interacts with Kipferl and discovered that Kipferl interacts with Rhino’s chromodomain (Fig. 6H; Fig. S6G). Kipferl did not interact with the Rhino chromoshadow domain, the Rhino hinge region, or the chromodomain of Su(var)2-5. To test these findings *in vivo*, we generated flies that expressed Rhino variants with artificial hinge or with a Gcn4 dimerization domain instead of the chromoshadow domain. Both Rhino^CSD-Gcn4^ and Rhino^art.-hinge^ failed to form extended Rhino domains like the wildtype protein, but were enriched at prominent Kipferl nucleation sites such as those in *cluster 80F* or upstream of the *headcase* gene (Fig. 6I). Flies that expressed a Rhino variant with the chromodomain of Su(var)2-5 had rudimentary ovaries, precluding a meaningful ChIP-seq analysis, further supporting a model where the chromodomain contributes more than just binding to methylated H3K9. Our combined data indicate that Kipferl recruits Rhino to chromatin via a direct contact between its 4^th^ ZnF and the Rhino chromodomain, and that functional Rhino is required for the extension and strengthening of Rhino/Kipferl domains.

### Kipferl is required for Rhino domains at diverse stand-alone transposon loci

The piRNA pathway’s primary role is to silence transposable elements. Given their sequence diversity, it was surprising to find that a DNA binding protein with affinity for a short nucleotide motif is required for Rhino’s chromatin occupancy and hence the determination of the ovarian piRNA pool. In fact, transposon sequences overall do not harbor more GRGGN motif occurrences than random genomic tiles (Fig. S7A). Instead, we observed a large spread of motif density among transposons (Fig. S7B). The GC-rich *gypsy8* and *Rt1a/b/c* elements harbored between 29 and 38 motives, while the AT-rich *Rsp* and *1.688* Satellites harbored none (Fig. 7A; Fig. S7C). Motif density per kilobase showed a significant correlation with Kipferl’s intrinsic ability to bind to transposon sequences (as measured by Kipferl ChIP-seq enrichment in *rhino* mutant ovaries; R = 0.64, p < 2.2e-16) (Fig. 7B). For most elements, the extent of Rhino-independent Kipferl enrichment was moreover directly correlated with their respective Rhino enrichment levels in wildtype ovaries (Fig. 7A, C). Exceptions were telomeric transposons and a few elements contained in clusters *42AB* or *38C*; these transposons were occupied by Rhino in wildtype ovaries despite no baseline Kipferl binding, and they maintained Rhino binding in *kipferl*-depleted ovaries, supporting the previous notion that Kipferl-independent chromatin recruitment mechanisms for Rhino likely exist (Fig. 7C; Fig. S7D). As the occurrence of the guanosine-rich Kipferl motif correlated with overall GC-content, enrichments for both Kipferl and Rhino on transposons also correlated with the elements GC-content (Fig. S7A, E, F). The correlation between GC-content and Rhino enrichment was abolished in ovaries depleted for Kipferl, indicating that Rhino’s preferential binding to GC-rich transposons is due to Kipferl-mediated recruitment and/or stabilization of Rhino on DNA sequences with Kipferl motifs (Fig. S7G).

**Fig. 7:**
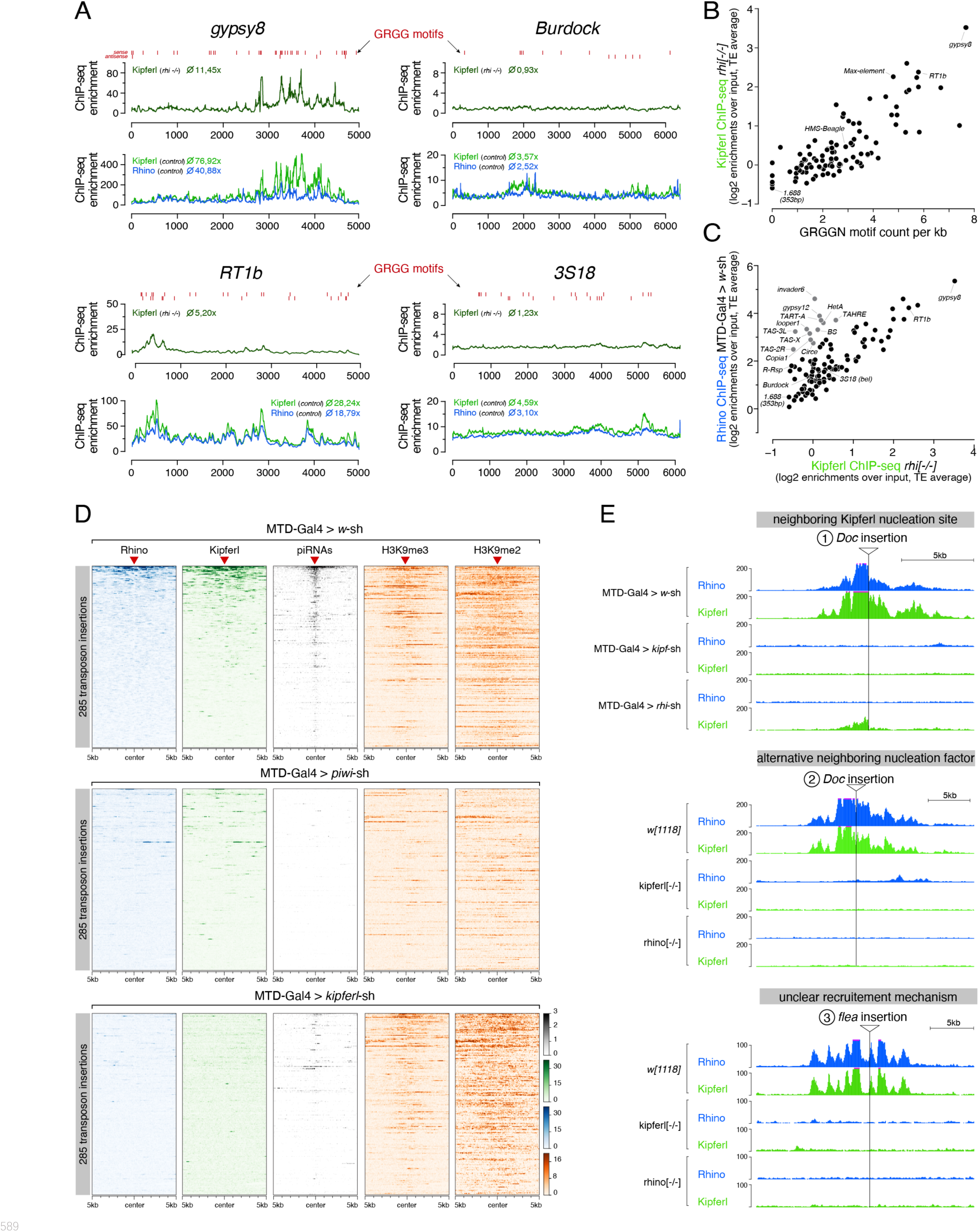
Kipferl is required for the establishment of Rhino domains at stand-alone transposon loci. **(A)** ChIP-seq enrichment profiles on consensus sequences of transposons with high (*gypsy8* and *Rt1b*) or low (*Burdock* and *3S18*) number of Kipferl DNA binding motifs per kb sequence. Indicated ChIP-seq signals are displayed as average enrichment over input in two (Kipferl) or three replicates (Rhino) of ovaries from *rhino* mutant (top tracks) or MTD-Gal4 > *w*-sh control ovaries (bottom tracks). Red bars indicate motif instances on the sense or antisense strand. Numbers indicate average enrichment across the entire element. **(B)** Scatter plot correlating the GRGGN motif count (normalized to element length) to the Rhino-independent Kipferl ChIP-seq enrichment for each transposon (ChIP-seq enrichments depict average of two independent experiments). **(C)** Scatter plot depicting the relation between wildtype Rhino ChIP-seq enrichments and Rhino-independent Kipferl ChIP-seq enrichments per transposon (average of two (Kipferl) or three (Rhino) independent experiments; elements indicated in grey are bound by Rhino in a largely Kipferl-independent manner, see also Fig.S7D). **(D)** Heat maps depicting indicated ChIP-seq signal in the genomic regions flanking 285 euchromatic stand-alone insertions (red triangles) of Rhino-dependent transposons with low Rhino-independent Kipferl binding (data sorted by the ChIP-seq signal detected for Rhino in MTD-Gal4 > *w*-sh ovaries). **(E)** UCSC genome browser tracks of stand-alone transposon insertions found in *MTD-Gal4* or *w[1118]* strains, depicting examples of different potential modes of Kipferl dependency. ChIP-seq signal is shown as coverage per million reads (genomic coordinates of individual panels listed in Table S3).

Many transposons did not show baseline Kipferl binding, yet their low-level Rhino enrichment in wildtype ovaries still depended on Kipferl. Moreover, several of these transposons (e.g. *Burdock*, *HMS-Beagle*, *3S18*) were strongly dependent on Rhino for their silencing, and were deregulated also upon loss of Kipferl, although to a weaker extent (Fig. 7A; Fig. S7H, I). The recent observation that flies mutant for the three piRNA clusters *38C*, *42AB*, and *20A* are fully fertile and do not show upregulation of these elements (Gebert et al., 2021) supports the proposal that stand-alone insertions likely act as independent Rhino domains (Mohn et al., 2014, Shpiz et al., 2014), providing piRNAs capable of targeting other insertions in *trans*. Considering that only about 20% of transposon insertions are transformed into Rhino domains (Akulenko et al., 2018), the average Rhino enrichment mapped to the consensus sequence would be low. We determined all stand-alone insertions of transposons lacking baseline Kipferl-binding in the MTD-Gal4 strain and displayed the levels of Rhino, Kipferl, H3K9me2/3, and piRNAs in the flanking genomic regions of these insertions (Fig. 7D). While nearly all insertions were embedded in a local H3K9me2/3 domain, Rhino and Kipferl were only enriched at a subset of these insertions. RNAi-mediated depletion of Piwi or Kipferl resulted in loss of Rhino and piRNAs at these insertions. However, while Piwi loss impaired local heterochromatin formation, Kipferl loss did not (Fig. 7D). We conclude that a combination of Kipferl and Piwi-dependent H3K9me2/3 is required to stabilize Rhino even on those transposons that do not contain strong Kipferl nucleation sites.

Based on manual inspection of individual transposon insertions in the different examined fly strains (illustrated in Fig. 7E) we propose that Kipferl supports Rhino at stand-alone transposon insertions in one of two ways: In some instances, we find that Rhino-bound insertions occurred nearby a genomic Kipferl binding site, which might be a critical factor to establish a local Rhino domain. In other instances, we find no nearby Kipferl binding site. Here, Kipferl likely acts by stabilizing Rhino which might have been recruited through alternative nucleation factors akin to Kipferl’s role at piRNA clusters *38C* and *42AB*, where Rhino remains chromatin bound in Kipferl mutant ovaries. Taken altogether, our data indicate that Kipferl acts as a recruitment factor for Rhino, and that it stabilizes the formation of extended Rhino domains, both at its own recruitment sites and at domains nucleated by alternative specificity factors.

## DISCUSSION

This study provides direct evidence that DNA sequence is an important determinant of how germline cells define the chromatin binding sites of the fast-evolving HP1 variant protein Rhino. The discovery of Kipferl, a DNA-binding zinc finger protein, that cooperates with the H3K9me3 chromatin mark and serves as Rhino guidance factor critically advances our understanding of how dual-strand piRNA source loci are specified.

By systematically comparing different *Drosophila melanogaster* strains, we find that Rhino domains in adult ovaries are largely identical between strains, yet highly diverse in their genomic location, underlying sequence content, and their piRNA output. Given the affinity of Rhino to the histone mark H3K9me3 (Mohn et al., 2014, Le Thomas et al., 2014, Yu et al., 2015), current models postulate that maternally inherited piRNAs define the cellular Rhino profile through epigenetic mechanisms during early embryogenesis (Mohn et al., 2014, Le Thomas et al., 2014, Shpiz et al., 2014, Akkouche et al., 2017). Indeed, we find that Rhino domains are invariably accompanied by local di- and/or tri-methylation of H3K9. However, our work uncovered several transposon-free Rhino domains, from where only negligible amounts of piRNAs are produced. These domains are difficult to explain by maternally deposited piRNAs as Rhino specifiers. Moreover, while Piwi depletion leads to a loss of Rhino from chromatin at stand-alone transposon insertions, where H3K9me2/3 is dependent on Piwi, most piRNA source loci, including the major piRNA clusters, are refractory to Rhino loss in the absence of Piwi (Mohn et al., 2014). This implied that other mechanisms must be in place to deposit Rhino at these sites. With Kipferl we present a factor acting upstream of Rhino that has a major impact on nearly all Rhino domains in ovaries, irrespective of their position, transposon content, or piRNA output. Importantly, Kipferl does not influence the H3K9 methylation status, but rather stabilizes Rhino at most of its genomic targets, where H3K9me2/3 is provided through parallel pathways. Thereby, Kipferl, with its specific DNA binding activity, serves as the first example of the long sought-after guidance cue(s) required for specifying Rhino’s binding profile within heterochromatin.

Our data indicate that Rhino binds to Kipferl via its chromodomain. Binding of client proteins has so far been assigned to the dimeric C-terminal chromoshadow domain of HP1 family proteins. By recruiting and/or stabilizing Rhino on chromatin via the chromodomain, Kipferl-binding would be compatible with the recruitment of downstream factors like Deadlock via Rhino’s chromoshadow domain (Yu et al., 2018). Our genetic data imply that DNA binding of Kipferl as well as H3K9me2/3 binding of Rhino are required for the formation of a stable Kipferl-Rhino complex on chromatin. Biochemical and structural work will be required to understand how the Kipferl-Rhino interaction evolved to be specific to the Rhino chromodomain and how it is compatible with simultaneous H3K9me2/3 binding. We were unable to generate Kipferl as a recombinant protein to study the Rhino-Kipferl interaction *in vitro*. It will further be interesting to investigate whether other chromatin readers utilize a similar mode of specialized recruitment for additional specificity. Known examples where ZnF proteins are involved in HP1 recruitment are the *Drosophila* HP1 variants HP1b and c, who rely on their cofactors Woc and Row for chromatin binding (Font-Burgada et al., 2008). In mouse embryonic stem cells, HP1beta/gamma bind to selected chromatin sites in a complex with the chromatin remodeler CHD4 and the activity-dependent neuroprotective protein (ADNP) (Ostapcuk et al., 2018). In both cases, interaction with ZnF proteins allows the H3K9me3-independent recruitment of HP1 variants to sites where their function has not been fully elucidated. This mode of recruitment is distinct from Kipferl’s mode of action, which synergizes with underlying H3K9me2/3 at its binding loci to recruit Rhino to chromatin.

With an N-terminal ZAD, Kipferl belongs to the largest group of ZnF proteins in insects, with 92 members in *Drosophila melanogaster* (Chung et al., 2007, Chung et al., 2002). The functions of the few characterized ZAD-ZnF proteins are diverse and range from transcriptional regulation to chromatin organization and heterochromatin biology. Many ZAD proteins are preferentially expressed in ovaries and during early embryogenesis (Shapiro-Kulnane et al., 2021), and some of them, despite having essential functions, are fast evolving (Kasinathan et al., 2020). It is therefore possible that additional ZAD-ZnF proteins function as guidance factors for Rhino or other HP1 proteins in the germline. If that were the case, the transposon-genome arms race might be a key driver of the diversification of ZAD-ZnF proteins. Diversification of ZnF containing genes is also observed for KRAB or SCAN domain zinc fingers, whose radiation in tetrapods is believed to be fueled by the transposon conflict (Bruno et al., 2019). While KRAB-ZnFs act directly as repressors of transposon expression and show fast evolution of their DNA binding specificity, signatures of positive selection are not concentrated on ZnF domains in ZAD proteins, indicating different evolutionary pressures acting on the two families of ZnF proteins.

An intriguing observation from our work is that Kipferl localizes to every Rhino domain in ovaries and is required for Rhino stability at most sites, but that its expression profile does not fully correlate with the expression of Rhino: Kipferl levels in ovarian germline stem cells and cystoblasts are very low, and the protein is absent in the testes. Kipferl’s ovary-specific expression is therefore a likely contributor for the pronounced differences in Rhino landscapes between ovaries and testes (Chen et al., 2021). At the same time, it implies the existence of additional Kipferl-like components or complementary molecular pathways that collaborate with H3K9me2/3 in testes or early developmental stages in ovaries where Rhino is functional in the absence of Kipferl.

At this point, we can only speculate about the processes that forced the evolution of a factor like Kipferl. The phenotype of *kipferl* mutants, however, holds important clues. In the absence of Kipferl, the piRNA profile in ovaries is strongly distorted. Most transposons exhibit reduced antisense piRNA levels, and for some this results in their de-repression and accumulation of transposon transcripts in the developing oocyte, potentially causative of the fertility defects of *kipferl* mutant females. One model would therefore be that Kipferl evolved as a dedicated Rhino specificity factor that targets the DNA sequence of some transposons, active in differentiating ovarian germline cells. While only few transposons contain strong Kipferl nucleation sites, we find that the strongest Kipferl binding sites genome-wide are within sequence fragments of the *gypsy8* and *DMRT1* family transposons. This indicates that the DNA motif recognized by Kipferl possibly originated from an ancient invasion of these now inactive elements, offering a potential evolutionary requirement for Kipferl as a direct recruitment factor for Rhino. The Rhino decoration of transposons lacking Kipferl nucleation sites might rely on sporadic genomic Kipferl binding sites which allow their specification as piRNA source loci as local chromatin state permits. The major piRNA *cluster 80F* would be a prominent example, where fragments of *gypsy8* and *Rt1A* and *B* insertions induce the Kipferl-dependent recruitment of Rhino and subsequent piRNA production also from neighboring transposons. In support of this, the entire *cluster 80F* is not a strong piRNA source in testes, where Kipferl is not expressed (Chen et al., 2021). Thus, an initially specific homing of Rhino to a subset of transposons via Kipferl might have evolved into a network of transposon insertions within heterochromatin and at selected sites in euchromatic chromosomal arms that co-depend on Kipferl. This system would offer the potential for robust Rhino recruitment, at the cost of occasional off-target Rhino domains, when Kipferl motifs and H3K9 methylation coincide at regions lacking transposon sequences. We can, however, also envision an alternative scenario, based on the second major phenotype in *kipferl* mutants: The dramatic re-localization of Rhino from hundreds of domains to Satellite arrays located within pericentromeric heterochromatin. Intriguingly, both the *Rsp* and the *1.688* Satellites are involved in genetic conflicts (Larracuente and Presgraves, 2012, Ferree and Barbash, 2009, Chen et al., 2021). The largest *1.688* satellite array, the X-chromosomal 359bp repeat, spans more than 10 Mbp on the pericentromeric X-chromosome and acts as a hybrid lethality locus in crosses between *D. melanogaster* and *D. simulans* (Ferree and Barbash, 2009, Chen et al., 2021). Females lacking this repeat, when mated to wildtype males, generate non-viable offspring while no defects occur in the reciprocal cross. This phenotype might stem from a requirement for maternally deposited piRNAs to prevent mitotic catastrophe caused by uncontrolled *1.688* repeats. The *Rsp* Satellite has been identified genetically as part of the Segregation Distorter system, a meiotic drive system in males (Larracuente and Presgraves, 2012). Both, *Rsp* and *1.688* Satellites give rise to abundant piRNAs in a Rhino-dependent manner in testes and ovaries (Wei et al., 2021, Chen et al., 2021). If piRNA production from *Rsp* and *1.688* Satellites is important to maintain control of *1.688* repeats or to suppress the *Distorter* locus during spermatogenesis, Satellites might have evolved to take advantage of the Rhino system and force piRNA production from their own loci. In this scenario, different genomic loci would compete for the cellular Rhino pool to recruit the Rhino-dependent transcription and export machinery. It is conceivable that proteins working in an analogous way to Kipferl are present at Satellites to sequester Rhino to these repeats. Intriguingly, Rhino is among the fastest evolving proteins in the fly genome (Vermaak et al., 2005). Previous models have postulated that the positive selection in Rhino is a consequence of the transposon-genome arms race (Yu et al., 2018). And even though we find no evidence that Kipferl is under positive selection, our findings point to the provocative possibility that selfish Satellite sequences might be among the central drivers behind Rhino’s fast evolution. We speculate that Kipferl might have evolved out of a necessity for a stabilizer of Rhino that allows it to bind its diverse genomic target loci and to avoid being sequestered by selfish Satellite repeats. While Kipferl’s affinity for guanosine-rich sequences optimally opposes the AT-rich satellite sequences, its low abundance together with the relatively simple DNA motif it recognizes might constitute an optimal level of promiscuous binding across the genome, allowing the targeting of diverse transposon families.

## ACKNOWLEDGMENTS

We thank the VBCF core facilities (NGS, VDRC) as well as the IMBA/GMI/IMP BioOptics facility for excellent support, and the IMBA Fly House for generating transgenic and CRISPR-edited fly lines. The Max Perutz Laboratories Monoclonal Antibody facility generated the CG2678 and Rhino hybridoma cell lines. Steven DeLuca shared the pUASz plasmid. Sarah Barnes helped with cloning of the Kipferl^ýZnF#4^ construct, Bernardo Almeida gave advice on motif analyses, and Maria Novatchkova supported bioinformatic analyses. We thank the Brennecke laboratory for help throughout this project and A. Larracuente, F. Mohn, J. Schnabl, and S. Barnes for comments on the manuscript.

## FUNDUNG STATEMENT

This research was funded by the Austrian Academy of Sciences, the European Community (ERC-2015- CoG-682181) and the Austrian Science Fund (F4303 and W1207). LB was funded by a Boehringer Ingelheim Fond PhD fellowship.

## AUTHOR CONTRIBUTIONS

LB performed all experiments, except establishment and analysis of the stable OSC line expressing Kipferl (SP). LB and DH performed the computational analyses. PD designed and generated all CRISPR engineered flies. JB supervised the project. LB and JB wrote the paper.

## DATA AND MATERIAL AVAILABILITY

Sequencing data sets have been deposited to the NCBI GEO archive (GSE202468). All fly strains generated for this study are available via the VDRC. Antibodies against Rhino and Kipferl are available upon request. All genome-wide sequencing data (ChIP-seq, small RNA-seq, RNA-seq) can be browsed at https://genome-euro.ucsc.edu/s/balisa/dm6_elife220506.

## MATERIALS AND METHODS

### Fly strains and husbandry

All fly stocks were kept at 25°C with 12h dark/light cycles. Fly strains with genotypes, identifiers, and original sources are listed in Table S4 and strains generated for this study are available from VDRC (http://stockcenter.vdrc.at/control/main). For ovary dissections, flies were aged for 2–6 days, and put on apple juice plates with fresh yeast paste for two days.

### Generation of transgenic and mutant fly strains

Frame-shift mutant alleles for *CG2678* and *rhino* were generated in isogenised *white* embryos after co-injection of plasmids pBS-Hsp70-Cas9 (Addgene #46294) and pU6-BbsI-chiRNA (Addgene #45946) modified to express sgRNAs. Whole locus *CG2678* deletion for RMCE was achieved following co-injection into ZH-2A(Act5C-Cas9) embryos (derived from Bloomington stock #58492) of plasmids pXZ13 (Zhang et al., 2014a) containing 1kb homology arms around a 3xP3-dsRed marker flanked by attP sites, together with pCFD4 (Addgene 49411) (Port et al., 2014) expressing two sgRNAs. sgRNA sequences are given in Table S5.

Fly strains harboring short hairpin RNA (shRNA) expression cassettes for germline knockdown were generated by cloning shRNA sequences into the Valium-20 vector (Ni et al., 2011) modified with a *white* selection marker (oligos: Table S4). Transgenic flies harboring GFP tagged wildtype or engineered Rhino constructs were generated via insertion of desired tag sequences under the control of the *rhino* promoter region and the *vasa* 3’UTR into the attP40 landing site (Markstein et al., 2008) in flies harboring a *rhino* frame shift mutation on the same chromosome. Overexpression constructs for *CG2678* were injected as pUASz plasmids (DeLuca and Spradling, 2018) containing the full intron-containing sequence of *CG2678* into the attP2 landing site (Groth et al., 2004). *CG2678* rescue constructs were introduced into the *CG2678* whole locus deletion flies through co-injection of pRVV578 plasmid (Addgene #108279) harboring the endogenous *CG2678* locus flanked by attB sites, with pBS130 (Addgene #26290) for expression of phiC31 integrase. Successful cassette exchange was monitored via loss of dsRed in eyes and the orientation of the inserted construct was verified by PCR. Rescue constructs harbor an internal 3xFLAG tag at residue S161, as neither amino- nor carboxy-terminal tagging of *CG2678* yielded fully functional protein. Exceptions are ZAD^delta^ and ZAD^GCN4^ constructs, which harbor an N-terminal 3xFLAG tag. Of note, we found that the CG2678 locus harbors an additional 25 amino acid exon, flanked by 109 and 31 nucleotides of intronic sequence up and downstream, respectively, in several laboratory fly strains. Comparison with protein sequences annotated for other *Drosophila* species confirm the presence of the additional in frame coding sequence, inserted between P347 and K347 of the annotated melanogaster *CG2678* protein. PCR analysis confirmed the absence of the respective DNA sequence from *iso1* genomic DNA, which served as the basis of the reference genome. The additional sequence is included in the *CG2678* overexpression construct. All RMCE rescue constructs harbor the reference genome locus.

### Generation of endogenous knock-in fly strains

Generation of endogenously tagged lines for *HP1b* and *HP1c* was achieved through co-injections of pU6-BbsI-chiRNA together with pBS donor plasmids containing 1kb homology arms into embryos from vas-Cas9; attP2 flies or ZH-2A(Act5C-Cas9) embryos for *HP1b* and *HP1c*, respectively. sgRNA sequences are listed in Table S5.

### Antibody generation

Mouse monoclonal antibodies against His-tagged CG2678 (aa M2-K188) and His-tagged Rhino (full length) were generated by the Max Perutz Labs Antibody Facility. Antigens were cloned in pET-15b, transformed in BL21(DE3) *E. coli*, and purified using Ni-NTA resin (QIAGEN) according to standard protocols. Polyclonal antibodies were raised against a CG2678 peptide (aa R171-I190) at Eurogentec.

### Cell lines

*Drosophila* ovarian somatic cells (OSC) cells were cultured as previously described (Niki et al., 2006, Saito et al., 2009). Stable OSC lines expressing internally tagged CG2678 under control of the *ubi63E* promoter and an SV40 3’UTR were generated by integration into an RMCE landing site.

### Western Blot

5 pairs of ovaries were mechanically disrupted in lysis buffer (20 mM Tris pH 8.0, 1% Triton X-100, 2 mM MgCl_2_, Benzonase, protease inhibitors) using a plastic pestle. Protein concentration of whole ovary lysate was determined via Bradford assay to allow equal loading, and SDS-PAGE gel electrophoresis was performed according to standard procedures. Primary antibodies were incubated at 4°C overnight, secondary antibodies for 1 hr at RT and the blots were developed using ECL (BioRad). Antibodies and dilutions are listed in Table S6.

### RNA Fluorescent In Situ Hybridization

RNA FISH for piRNA clusters *42AB* and *38C*, as well as *HMS-Beagle*, *Max*, *diver*, and *3S18* transposons was performed using Stellaris probes (Biosearch Technologies). Probe sequences are listed in Table S7. RNA FISH for *1.688* and *Rsp* Satellites was performed using a single fluorescent oligo or an in-house labelled probe set of 48 oligos, respectively (Wei et al., 2021, Gaspar et al., 2017). FISH was performed according to the manufacturers protocol with slight modification. 5 ovaries were dissected into ice-cold PBS, fixed at room temperature for 20 min (4% formaldehyde, 0.3% Triton X-100 in PBS), washed 3 times 5 min at RT (PBS containing 0.3% Triton X-100) and incubated at 4°C overnight in 70% EtOH to enhance permeabilization. Prior to hybridization, ovaries were rehydrated for 5 min in wash buffer (10% formamide in 2x SSC). Hybridization was done in 50μl hybridization buffer (100 mg/ml dextran sulfate and 10% formamide in 2X SSC) overnight at 37°C using 0.5 μl Stellaris and *Rsp* FISH probe per sample and a final concentration of 100 nM for the *1.688* FISH oligo. Samples were rinsed 2 times in wash buffer and then washed in wash buffer 2 times for 30 min at 37°C. Ovaries were counterstained for DNA (DAPI 1:5000 in 2x SSC) for 5 min at RT and washed 2 times 5 min with 2x SSC. Finally, ovaries were mounted on microscopy slides using DAKO mounting medium (Agilent) and equalized at RT for at least 24 hr prior to imaging on a Zeiss LSM 880 inverted Airyscan microscope.

### Immunofluorescence staining of ovaries and testes

5-10 ovary pairs or testes were dissected into ice cold PBS and subsequently incubated in fixation solution (4% formaldehyde, 0.3% Triton X-100, 1x PBS) for 20 minutes at room temperature with rotation. Fixed ovaries were washed 3x 5 minutes in PBX (0.3% Triton X-100, 1x PBS) and blocked with BBX (0.1% BSA, 0.3% Triton X-100, 1x PBS) for 30 minutes, all at room temperature with rotation. Primary antibody incubation was performed by incubation at 4°C overnight with antibodies diluted in BBX followed by three 5-minute washes in PBX. Ovaries were then incubated overnight at 4°C with fluorophore-coupled secondary antibodies, washed three times in PBX including DAPI in the first wash to stain DNA (1:50,000 dilution). The final wash buffer was carefully removed before addition of ∼40 μL DAKO mounting to each sample. The samples were imaged on a Zeiss LSM-880 Axio Imager confocal-microscope and image processing was done using FIJI/ImageJ (Schindelin et al., 2012). All relevant antibodies and dilutions are listed in the Table S6.

### Yeast Two-Hybrid Assay

#### Screening for Rhino interactors

Yeast two-hybrid screening for Rhino interactors was performed by Hybrigenics Services, S.A.S., Paris, France (https://www.hybrigenics-services.com). The coding sequence for full length Rhino was cloned into pB27 (derived from pBTM116) (Vojtek and Hollenberg, 1995) as an N-terminal fusion to LexA (LexA-Rhi). The construct was sequence verified and used as a bait to screen against a random-primed *Drosophila* ovary cDNA library constructed into pP6 (derived from pGADGH (Bartel et al., 1993). A mating approach with YHGX13 (Y187 ade2-101::loxP-kanMX-loxP, matα) and L40ΔGal4 (matA) yeast strains was used to screen 65 million interactions as previously described (Fromont-Racine et al., 1997). 173 colonies were selected on a medium lacking tryptophan, leucine, and histidine. The prey fragments of the positive clones were amplified by PCR and sequenced at their 5′ and 3′ junctions. The resulting sequences were used to identify the corresponding interacting proteins in the GenBank database (NCBI).

#### Validation and interaction mapping

Yeast strains were grown in YPD or SC selective medium at 30°C. pOAD and pOBD used as backbone for cloning were described in (Miller and Stagljar, 2004) (see Table S8 for yeast strains used in this study). Direct protein interactions were probed as described (Miller and Stagljar, 2004). In brief, assayed proteins were fused to the activation domain (AD) and DNA-binding domain (DBD) of the Gal4 transcription factor and transformed into yeast strains PJ694A (AD) and PJ694α (DBD). Individually transformed colonies were selected, picked and mated. Interactions were detected upon spotting of a dilution series of mated yeast on selective (-LTH) plates. Parallel plating on non-selective (-LT) plates controlled for presence of both plasmids.

### Scoring of embryo hatching rates

To measure female fertility, 10 virgin females were collected and aged for 2-3 days with at least 24h of mating with three *w^1118^* males. The hatching rate of fertilized eggs laid onto apple juice plates within a period of 4-7 hours was determined 30 hours after egg laying (25 degrees), as percentage of hatched eggs from total. Plates with less than 50 eggs were disregarded in the analysis. Wildtype females were included in every experiment as control.

### Definition and Curation of 1 kb Genomic Windows

The four assembled chromosomes of the *Drosophila melanogaster* genome (dm6 assembly) were split into non-overlapping 1-kb tiles. The tiles were annotated by intersection with genomic annotations for piRNA clusters. Tile mappability was determined by intersection with genomic blocks of continuous mappability using bedtools coverage. Tiles with mappability below 25% were excluded from all analyses (2761 1-kb tiles). Further exclusion criteria included a more than three-fold deviation from median values for representative input libraries for either of the three wildtype genotypes used in this study (*w^1118^*, *MTD- Gal4* > *w*-sh, *iso1*; affecting 18,268 1-kb tiles), as well as tiles showing strong residual Rhino or CG2678 signal in ChIP-seq libraries prepared from the respective knock out ovaries (20 and 495 tiles, respectively).

### Heterochromatin and Euchromatin Definitions Used in This Study

We used ovary H3K9me3 ChIP-seq data to define the extent of pericentromeric heterochromatin and euchromatic chromosome arms. The heavily H3K9me3 covered pericentric regions of the assembled chromosomes, as well as the entire chromosome 4 were classified as heterochromatic, while the rest was annotated as euchromatic. Detailed coordinates can be found in Table S1. Small genome contigs not assembled into the four major chromosomes were excluded from all analyses.

### ChIP-Seq

ChIP was performed as previously described (Lee et al., 2006). In brief, 150 μl of ovaries were dissected into ice-cold PBS, crosslinked with 1.8% formaldehyde in PBS for 10 min at room temperature, quenched with Glycine, rinsed in PBS and flash frozen in liquid nitrogen after removing all PBS. Frozen ovaries were disrupted in PBS using a dounce homogenizer, centrifuged at low speed and the pellet was resuspended in lysis buffer. For ChIP from OSCs 5-10 million cells were crosslinked, quenched, and lysed. Sonication (Bioruptor) resulted in DNA fragment sizes of 200–800 bp. Immunoprecipitation with specific antibodies was done overnight at 4°C in 350–700 μl total volume using 1/3 to 1/4 of chromatin per ChIP. Then, 40 μl Dynabeads (equal mixture of Protein G and A, Invitrogen) were added and incubated for 1 hr at 4°. After multiple washes, immuno-precipitated protein-DNA complexes were eluted with 1% SDS, treated with RNAse-A, decrosslinked overnight at 65°C, and proteins were digested with proteinase K before clean-up using ChIP DNA Clean & Concentrator columns (Zymo Research). Barcoded libraries were prepared according to manufacturer’s instructions using the NEBNext Ultra II DNA Library Prep Kit for Illumina (NEB), and sequenced on a HiSeqV4, NextSeq550, or NovaSeqSP (Illumina).

### RNA-Seq

Strand-specific RNA seq was performed as described previously (Zhang et al., 2012b). In brief, total RNA was extracted from 5–10 ovaries from 7 day old flies using Trizol (Invitrogen). Total RNA was purified using RNAeasy columns (QIAGEN). Six micrograms of total RNA were subjected to polyA selection and subsequent fragmentation, reverse transcription, and library preparation according to manufacturer’s instructions using the NEBNext Ultra DNA Library Prep Kit for Illumina (NEB) for sequencing on an Illumina NovaSeqSP instrument.

### Small RNA-Seq

Small RNA cloning was performed as described in (Grentzinger et al., 2020). In brief, ovaries were lysed and Argonaute- sRNA complexes were isolated using TraPR ion exchange spin columns. sRNAs were subsequently purified using Trizol and subjected to ligations of 3′ and 5′ barcoded adapters containing 4 random nucleotides at the ends to reduce ligation biases, reverse transcribed, PCR amplified, and sequenced on an Illumina NextSeq550 instrument.

### Computational Analysis

#### ChIP-Seq Analysis

ChIP-seq reads were trimmed to remove the adaptor sequences and to adjust all reads to 50 bp irrespective of sequencing mode. Reads were mapped to the dm6 genome using Bowtie (version.1.3.0, settings: -f -v 3 -a --best --strata --sam), allowing up to three mismatches. Genome unique reads were mapped to 1-kb tiles and a pseudocount of 1 was added after normalization to library depth, before enrichment over input values were determined. Each ChIP-seq sample was adjusted with a correction factor determined from median input levels and median background levels to reach median background enrichment of 1 to correct for unequal ChIP efficiency. To classify genomic regions into Rhino domains and non-Rhino domains, we used a binary cutoff of 4-fold enrichment calculated from two independent replicate experiments of the relevant wildtype genotypes. This cutoff corresponds to a p-value of < 0.05. Kipferl-only 1-kb tiles were those that had no Rhino enrichment (below 4-fold) and that were significantly enriched in Kipferl over Rhino (Z-score = 3).

#### ChIP-seq peak calling

We used MACS2 (Zhang et al., 2008) with --broad --broad-cutoff 0.1 for Kipferl and Rhino in wildtype ovaries due to the broad extent of Rhino/Kipferl domains. The ‘narrow peak’ setting was used for the remaining experiments. Peaks mapping to genomic contigs outside the four main chromosomes were discarded, and peaks were filtered for a score of 50 (broad peaks; p<10^-5^) and 30 (narrow peaks; p<10^-3^). Kipferl-Rhino shared versus Kipferl-only peaks were distinguished by intersection of broad peaks called for the two proteins in two independent replicate experiments of *w^1118^*ovaries using bedtools intersect with -u -f 0.75 for shared domains and -v for Kipferl-only domains. Rhino-independent Kipferl peaks that were detected independently in two replicate experiments were grouped into heterochromatic and euchromatic by intersection with heterochromatin coordinates outlined above. Kipferl DNA binding motifs were recovered from the top ∼3000 summits of Rhino-independent Kipferl peaks (achieved through a score cutoff of 7 on summits) using HOMER (Heinz et al., 2010). Heatmaps display one representative replicate and were produced through deeptools (Ramirez et al., 2016).

#### ChIP-seq analysis on transposon consensus sequences

Genome mapping reads longer than 23 nucleotides were mapped to TE consensus sequences (Table S9) using bowtie (v.1.3.0; settings: -f -v 3 - a --best --strata --sam) allowing up to 3 mismatches. Reads mapping to multiple elements were assigned to the best mapping position. Reads mapping to multiple positions were randomly distributed. Library depth normalized ChIP and input reads, respectively, were averaged over all nucleotide positions of each element to give one value per element. ChIP-seq enrichment was calculated after adding a pseudo count of 1 and adjusted using sample-specific correction factors determined from background 1 kb tiles to reach median background enrichments of 1. Corrected per-base enrichment was calculated for TE ChIP-seq profiles.

#### Motif instances

Occurrences of Kipferl DNA binding motifs were determined using PWMScan (ccg.epfl.ch/pwmtools /pwmscan.php) on the dm6 genome and on TE consensus sequences. For display in heatmaps, cumulative motif counts on both genomic strands were intersected with non-overlapping 100 bp windows.

#### smallRNA-Seq Analysis

Raw reads were trimmed for linker sequences and the 4 random nucleotides flanking the small RNA before mapping to the *Drosophila melanogaster* genome (dm6), using Bowtie (version.1.3.0, settings: -f -v 3 -a --best --strata --sam) with 0 mismatch allowed. Genome mapping reads were intersected with Flybase genome annotations (r6.40) using Bedtools to allow the removal of reads mapping to rRNA, tRNA, snRNA, snoRNA loci and the mitochondrial genome. For TE mappings, all genome mappers were used allowing no mismatches. Reads mapping to multiple elements were assigned to the best match. Reads mapping equally well to multiple positions were randomly distributed. Libraries were normalized to 1 Mio miRNA reads. For the 1 kb window analysis, a pseudocount of 1 was added after normalization to library depth and correction for the mappability of the respective 1-kb tile. Tiles with fewer than 10 mapping piRNAs were disregarded for sRNA analysis to avoid distortion due to very low abundant piRNAs. For calculation of piRNAs mapping to TEs, sense and antisense piRNAs were kept separate, and counts were normalized to TE length. For classification of tiles and transposons into somatic, Rhino-independent, and Rhino-dependent source loci, the soma index was determined as the log2 ratio of somatic (Piwi-IP in *piwi* GLKD, normalized to library depth) and germline (GL-Piwi IP, normalized to library depth) piRNAs mapping to each tile or TE (Mohn et al., 2014). Classification by Kipferl-dependency of TEs was achieved by a binary cutoff of at a 2-fold reduction in antisense piRNA levels in *kipferl* knock down compared to control.

#### RNA-Seq Analysis

For the RNAseq analysis, genome matching reads (STAR v2.7.10a ; settings: -- outSAMmode NoQS --readFilesCommand cat --alignEndsType Local --twopassMode Basic -- outReadsUnmapped Fastx --outMultimapperOrder Random --outSAMtype SAM -- outFilterMultimapNmax 1000 --winAnchorMultimapNmax 2000 --outFilterMismatchNmax 3 -- seedSearchStartLmax 30 --alignSoftClipAtReferenceEnds No --outFilterType BySJout -- alignSJoverhangMin 15 --alignSJDBoverhangMin 1) were randomized in order and quantified using Salmon (v.1.7.0; settings: --dumpEqWeights --seqBias --gcBias --useVBOpt --numBootstraps 100 -l SF --incompatPrior 0.0 --validateMappings). For the analysis we used the FlyBase transcriptome (r6.40) which has been masked for sequences similar to transposons. To include both strands of transposons in the analysis, TE-consensus sequences were added to the FlyBase transcriptome in sense and antisense orientation. For gene expression visualization Salmon results were further processed to GeTMM values using edgeR (v3.34.0). For differential gene expression analysis Salmon results were processed using DeSeq2 (v1.32.0).

#### TE Insertion Calling

Euchromatic TE insertions were extracted from insertions called previously (Mohn et al., 2014), and nearby insertions were merged using bedtools merge -d 100, as these mostly corresponded to the same insertions called at slightly different positions. Further, all insertions overlapping UCSC repeat masker track annotations were discarded. For analysis of non-Kipferl nucleation site containing, but Rhino-dependent TEs, the resulting list was subsequently filtered for elements with no Kipferl enrichment in *rhino* knock out ovaries and at least 10-fold difference in piRNA levels between control and *rhino* MTD-Gal4 mediated germline knock down. This retrieved 285 euchromatic solo TE insertion sites in the genome of our experimental MTD-Gal4 strains (Table S10).

---

**Fig. S1.**
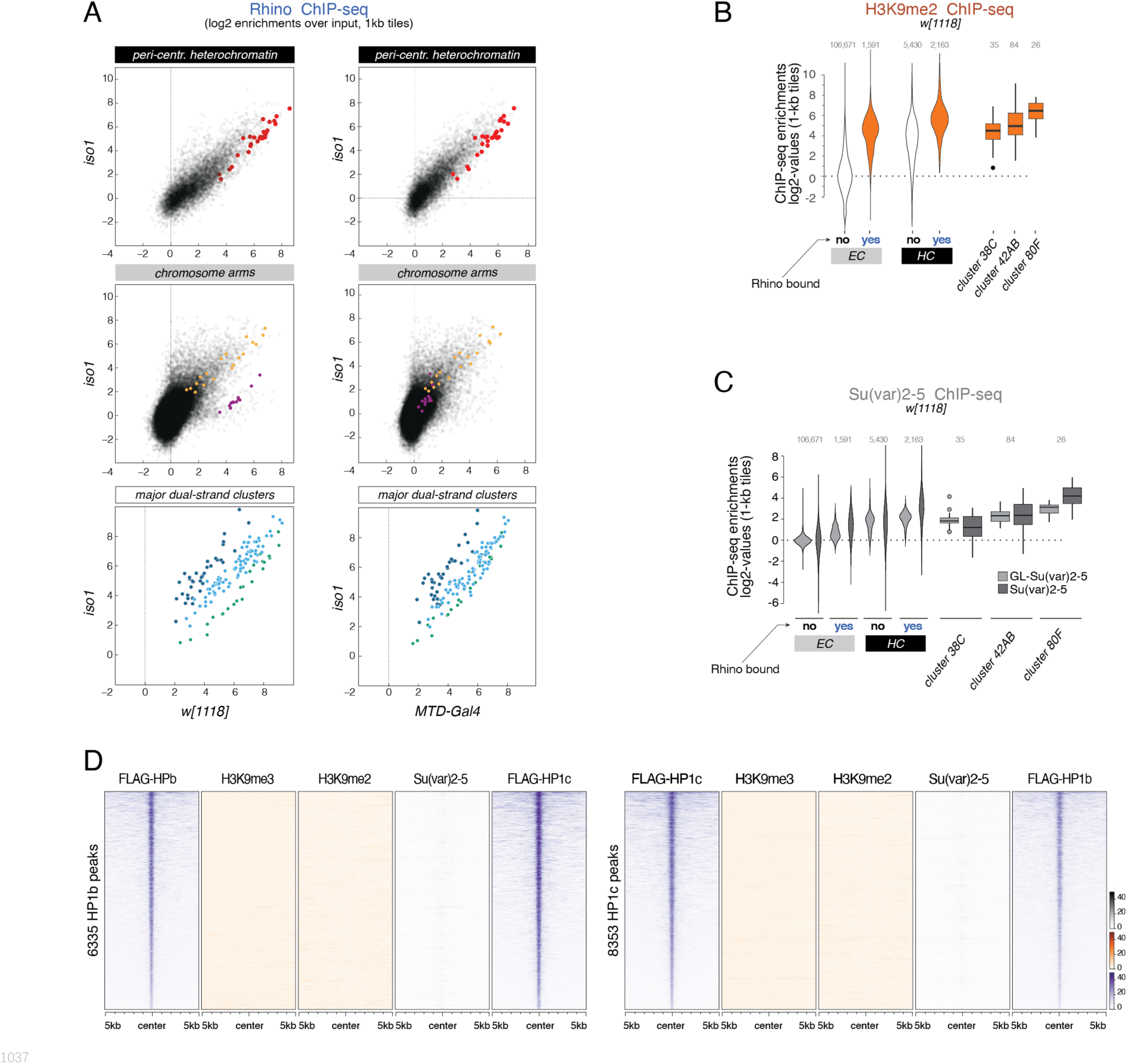
(related to Figure 1) **(A)** Scatter plots contrasting average log2 ChIP-seq enrichment for Rhino in *w[1118]* (n=2) or MTD-Gal4 (n=3) versus *iso1* (n=2) ovaries. Depicted are 1-kb tiles separated into pericentromeric heterochromatin and chromosomal arms. piRNA clusters *38C*, *42AB*, and *80F* are excluded from the categorization into heterochromatin and euchromatin and are depicted separately. Indicated colored 1-kb tiles correspond to example loci in Fig. 1F. **(B, C)** Violin plots showing the average log2-fold ChIP-seq enrichment of endogenous and germline specific Su(var)2-5 (transgene expressed under the control of the *rhino* promoter) and H3K9me2 (C) over input on 1-kb tiles. Tiles are grouped into Rhino-bound and non-Rhino-bound analogous to Fig. 1D and E with Rhino-dependent piRNA clusters *38C*, *42AB*, and *80F* depicted separately for comparison. **(D)** Heat map centered on HP1b (left) or HP1c (right) peaks, respectively, depicting the indicated ChIP-seq signal (n=1) in ovaries from the *w[1118]* strain.

**Fig. S2.**
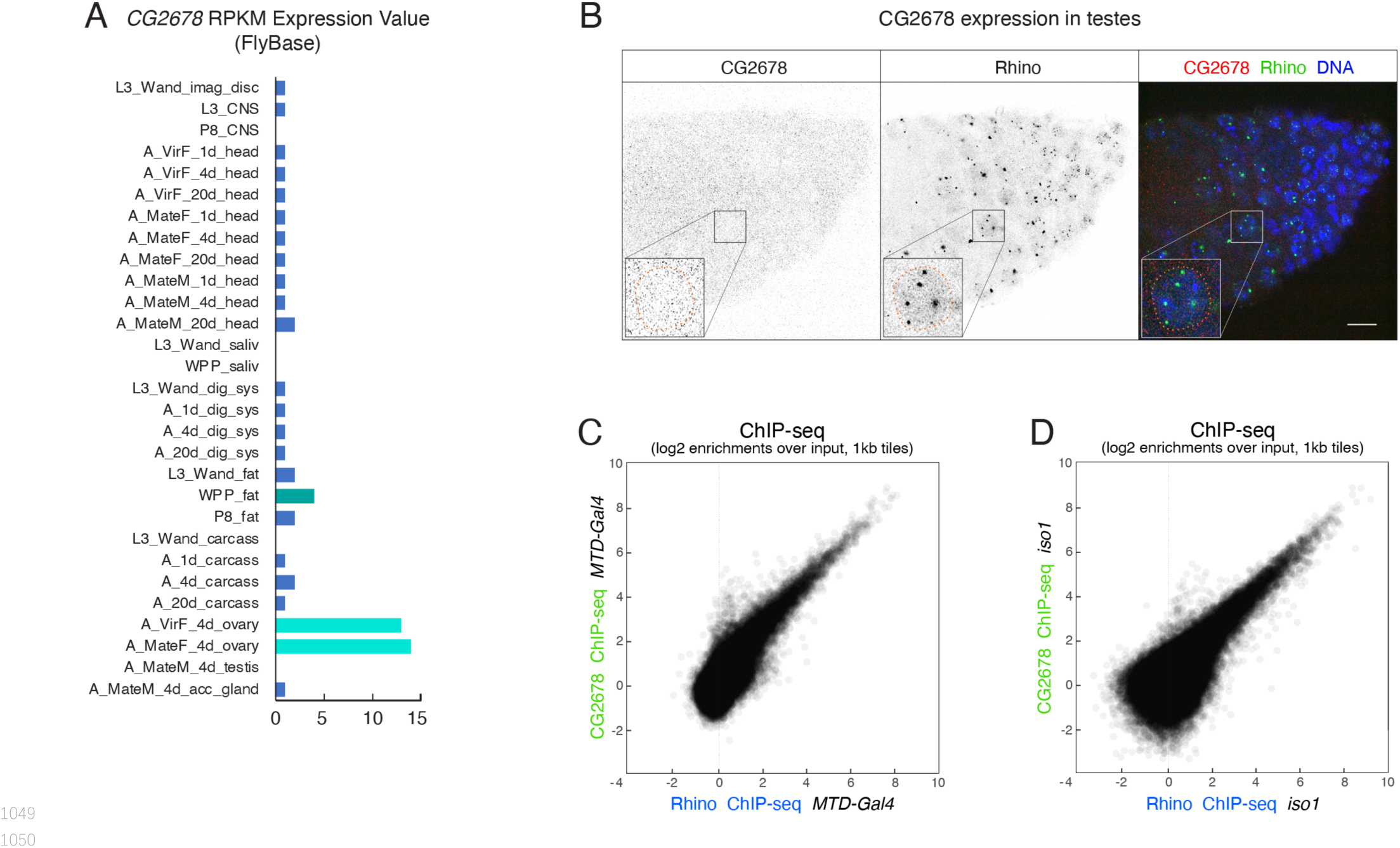
(related to Figure 2) **(A)** Bar graph indicating the expression pattern of *CG2678* (Flybase modENCODE Anatomy RNA-Seq; L: larva; P: pupa; A: adult; M: male; F: female) **(B)** Confocal image of anterior tip of a testes stained for endogenous CG2678 (left, red) and Rhino (middle, green) with enlarged view of one nucleus (dotted line: nuclear outline based on DAPI; scale bar: 5 µm). **(C, D)** Scatter plots depicting the correlation of log2-fold Rhino- versus CG2678 ChIP-seq enrichments in MTD-Gal4 (C; n=2) or *iso1* ovaries (D; n=1).

**Fig. S3.**
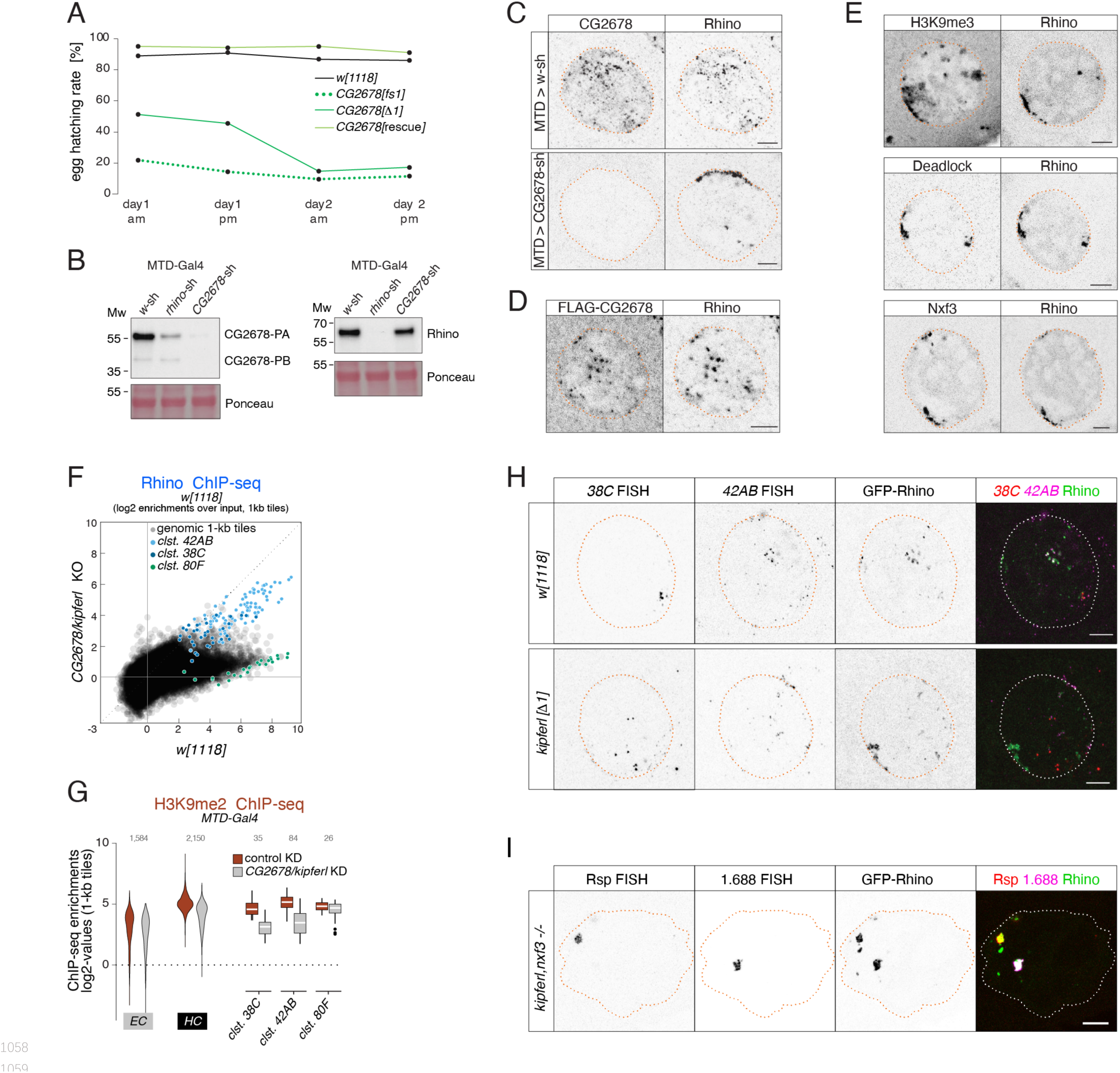
(related to Figure 3) **(A)** Time-resolved egg hatching rates for eggs laid by *w[1118]* control females in comparison to females carrying a *CG2678* frame shift (fs1), locus deletion (Δ1), or tagged rescue construct instead of the CG2678 locus, respectively (AM, PM indicates egg laying time). **(B)** Confocal images showing the localization of GFP-Rhino with H3K9me3 (top), Deadlock (middle), or Nxf3 (bottom) in *CG2678* null mutant nurse cells (scale bar: 5 µm). **(C)** Confocal images showing immunofluorescence signal for CG2678 and Rhino in nurse cells of MTD-Gal4 driven control or *CG2678* knockdown ovaries (scale bar: 5 µm). **(D)** Western blots showing CG2678 or Rhino protein levels in ovarian lysate from indicated genotypes (Ponceau staining served as loading control). **(E)** Confocal images showing FLAG-tagged CG2678 and Rhino signal in nurse cells expressing only an internally FLAG-tagged CG2678 (see Fig. 2B for location of FLAG tag; scale bar: 5 µm). **(F)** Scatter plot of genomic 1-kb tiles contrasting average log2-fold enrichment for Rhino ChIP-seq signal in *CG2678* knock out versus *w[1118]* control ovaries (three and two replicate experiments, respectively). **(G)** Violin plots showing average log2-fold H3K9me3 ChIP-seq enrichment for 1-kb tiles in MTD-Gal4 background in heterochromatin (HC) and along chromatin arms (EC) contrasting control to RNAi-mediated *CG2678/kipferl* knockdown. Rhino- dependent piRNA clusters *38C*, *42AB*, and *80F* are shown separately. **(H)** Confocal images showing the localization of piRNA cluster RNA via RNA FISH in respect to GFP-Rhino in nurse cells of *w[1118]* or *CG2678/kipferl* locus deletion flies (scale bar: 5 µm). **(I)** Confocal images showing *Rsp* and 1.*688* Satellite RNA FISH signal in respect to GFP-Rhino in nurse cells of *CG2678/kipferl,nxf3* double mutant flies (scale bar: 5 µm).

**Fig. S4.**
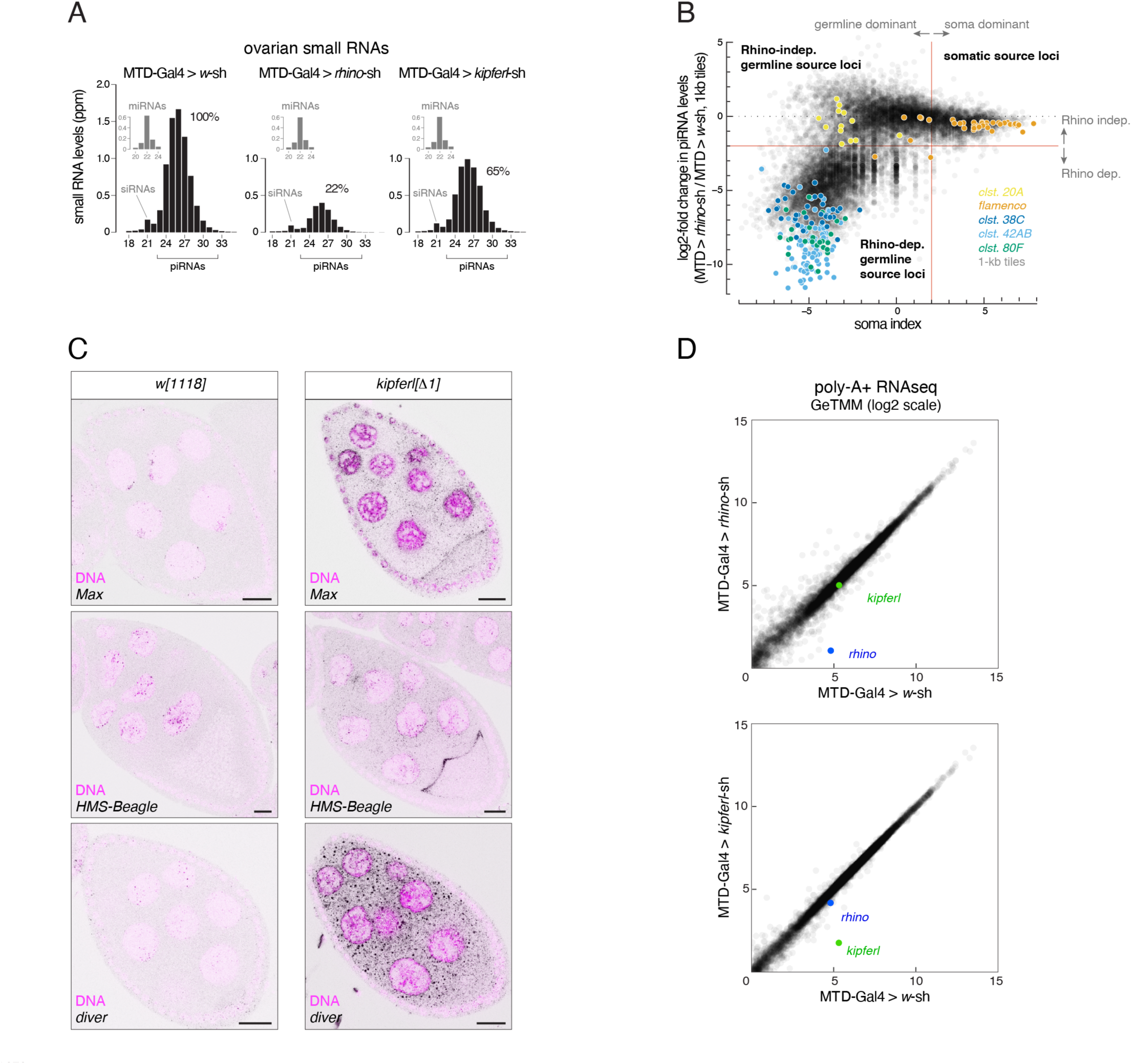
(related to Figure 4) **(A)** Size profiles of Argonaute-bound small RNAs (miRNA-normalized) isolated from ovaries of indicated genotypes. microRNA reads are shown separately (grey bars). Percentages represent total piRNA levels (23-33nt) relative to control (100%). **(B)** Classification of genomic 1-kb tiles into somatic (e.g. *flamenco*), Rhino-independent (e.g. cluster *20A*) and Rhino-dependent germline source loci (e.g. clusters *38C, 42AB, 80F*). Binary cutoffs are applied at a soma index of 2 do distinguish somatic from germline source loci (see Methods section) and a 4-fold reduction in piRNAs upon *rhino* depletion to determine dependency on Rhino. **(C)** Confocal images showing RNA FISH signal of transcripts for the indicated transposons in *w[1118]* and *kipferl* whole locus deletion egg chambers (scale bar: 20 µm; purple signal: DAPI). **(D)** Scatter plots contrasting RNAseq signal (poly-A plus) for expressed genes in indicated genotypes. Values are displayed as Gene length corrected trimmed mean of M-values (GeTMM); blue dot: *rhino*. Green dot: *kipferl*).

**Fig. S5.**
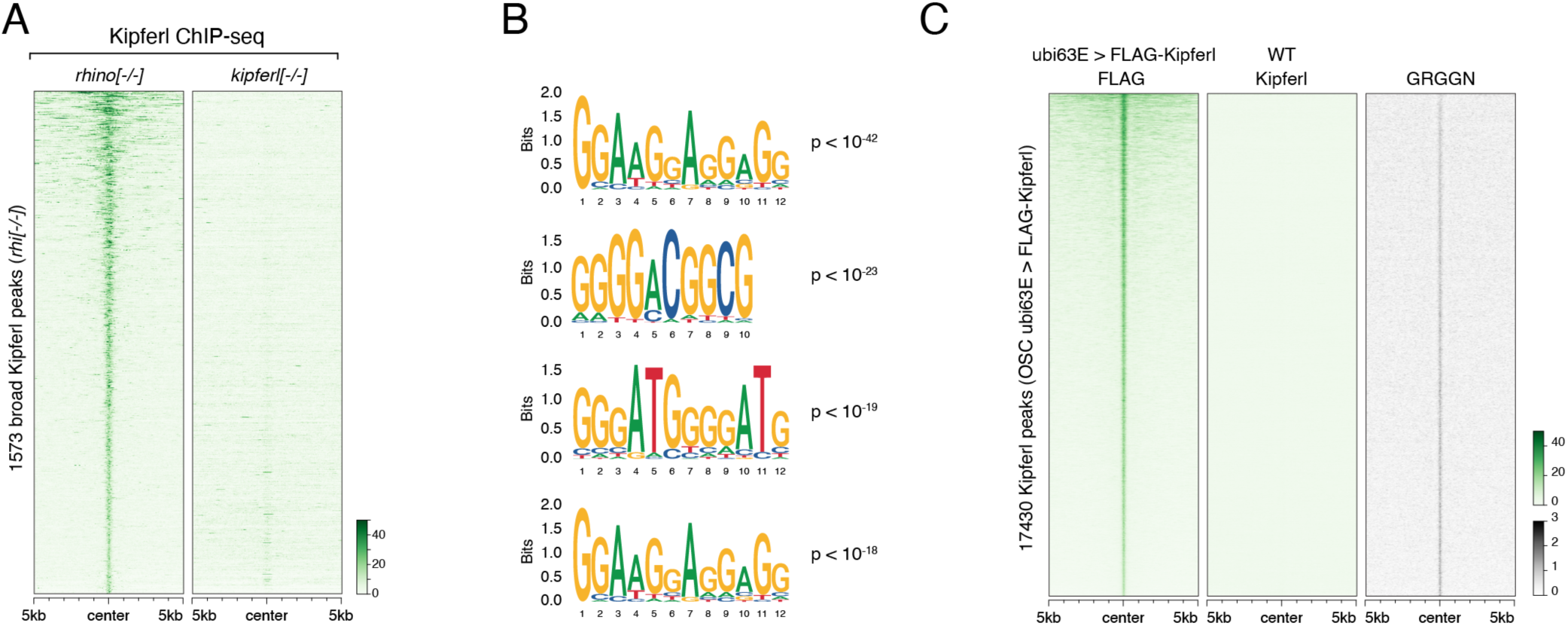
(related to Figure 5) **(A)** Heat map centered on broad Kipferl peaks detected in *rhino* mutant ovaries. Kipferl ChIP signal is depicted for *rhino* and *kipferl* mutant ovaries, respectively, as coverage per million reads. **(B)** Position weight matrices for highest ranking additional motifs recovered in the top 3,000 narrow Kipferl peaks present in two independent replicates of *rhino* mutant ovaries (p-values indicate significance of motif enrichment over sequence matched control regions). **(C)** Heat map centered on Kipferl peaks detected via FLAG ChIP-seq in cultured ovarian somatic stem cells (OSCs) stably expressing FLAG-tagged Kipferl under control of the *ubi63E* promoter. GRGGN motif count is given as count per non-overlapping genomic 100 bp window.

**Fig. S6.**
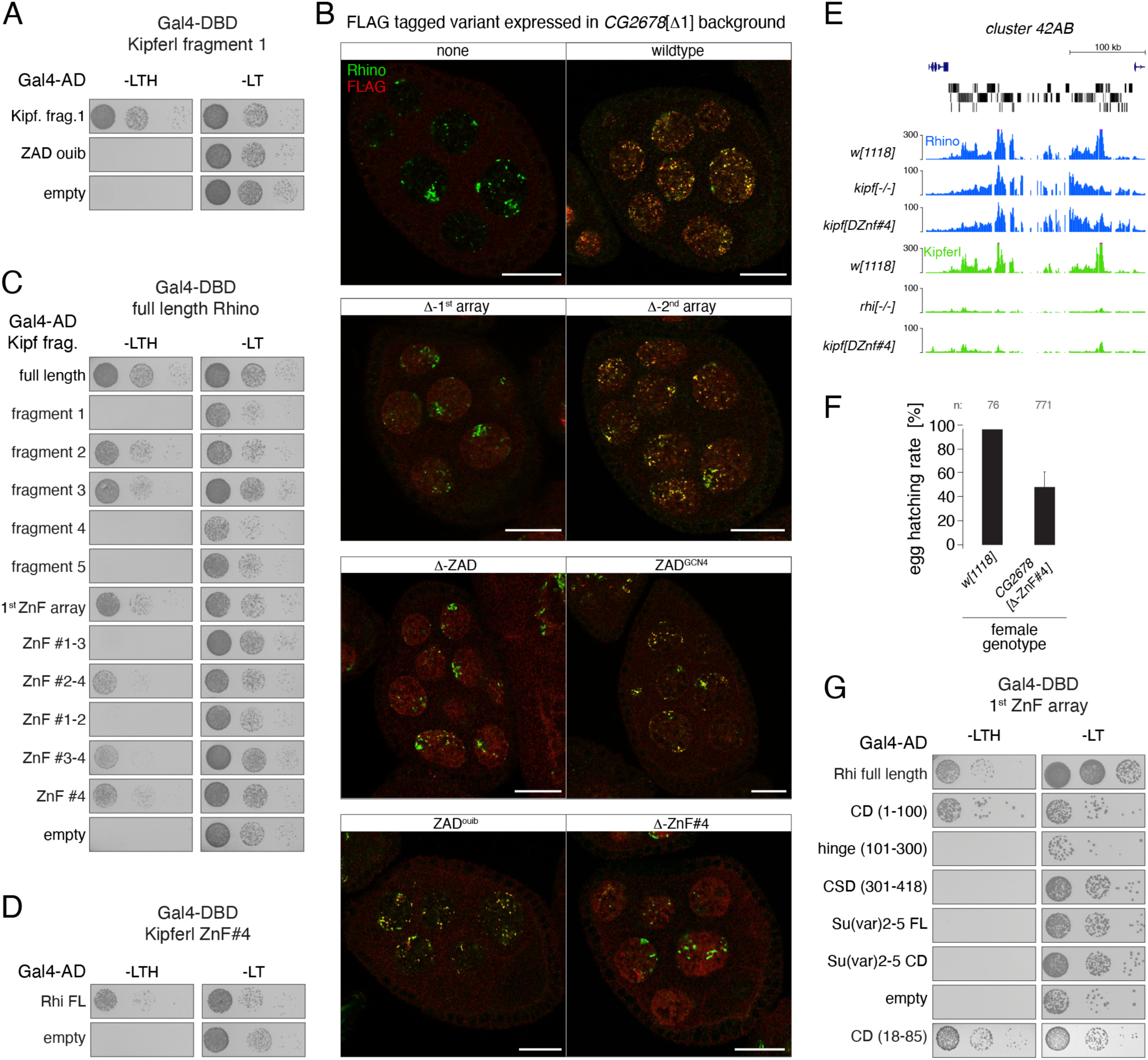
(related to Figure 6) **(A, C, D)** Yeast two hybrid assay on selective (left) and nonselective medium (right). AD: Gal4 activation domain; DBD: Gal4 DNA binding domain. Fragment identity is indicated in Fig. 6C. **(B)** Confocal images showing entire egg chambers with immunofluorescence stainings for Rhino (green) and indicated FLAG-tagged Kipferl variants (red) (scale bars: 20 µm). **(E)** USCS genome browser tracks showing indicated ChIP-seq signal as coverage per million sequenced reads in indicated genotypes at piRNA cluster *42AB* (genomic coordinates listed in Table S3). **(F)** Bar graph indicating average embryo hatching rate of eggs laid by *w[1118]* control females in comparison to females harboring a deletion of Kipferl’s ZnF #4. Data represents average of three technical replicates for Kipfer- ΔZnF#4 and one replicate for *w[1118]*. Total number of eggs laid is indicated as n. **(G)** Yeast two hybrid assay on selective (left) and nonselective medium (right). FL: full length. AD: Gal4 activation domain; DBD: Gal4 DNA binding domain. Fragment identity is indicated in Fig. 6H.

**Fig. S7.**
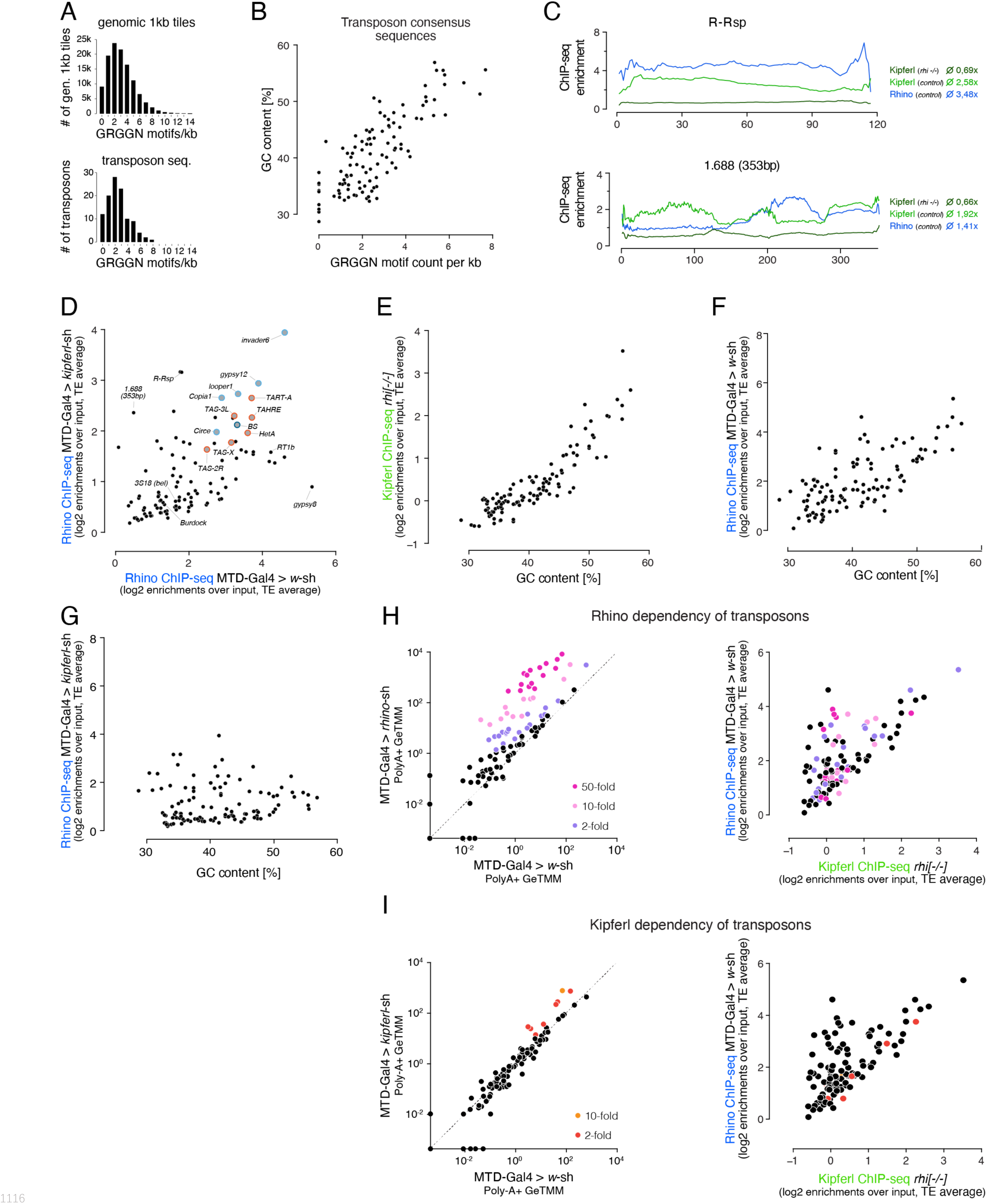
(related to Figure 7) **(A)** Bar graphs showing the distribution of motif frequency in all genomic 1-kb tiles (top) and in transposon consensus sequences (bottom). **(B)** Scatter plot of overall G/C content per transposon sequence versus the number of GRGGN motifs normalized to transposon length. **(C)** ChIP-seq enrichment profiles (over input) along the length of representative members of the *Rsp* and *1.688* family of Satellite repeats (both repeats lack Kipferl GRGGN motifs). Indicated ChIP-seq signals are displayed as average enrichment over input in two (Kipferl) or three replicates (Rhino) of ovaries from *rhino* mutant or control (MTD-Gal4 > *w*-sh) ovaries. Average enrichment across the entire element is indicated for each ChIP. **(D)** Scatter plot contrasting Rhino ChIP-seq enrichment (over input) on transposon sequences in control versus *kipferl*-depleted ovaries. Elements bound by Rhino independently of Kipferl (see Fig. 7C) are enlarged and highlighted in grey (those elements with insertions in piRNA clusters *42AB* and *38C* are additionally indicated in light and dark blue, respectively, and telomeric elements are indicated in red). **(E)** Scatter plot comparing GRGGN motif count per kb transposon sequence to average Rhino ChIP-seq enrichment in three replicate experiments on MTD-Gal4 > *w*-sh ovaries. (**F, G, H)** Scatter plots correlating the overall G/C nucleotide content per transposon to the ChIP-seq enrichment detected for Kipferl in *rhino* knock out ovaries (F) and for Rhino in wildtype (G) or *kipferl*-depleted ovaries. **(I, J)** Left: Classification of transposons into different grades of Rhino (I) and Kipferl (H) dependency based on poly-adenylated sense transcripts detected in wildtype control ovaries versus *rhino* or *kipferl*-depleted ovaries. Values are displayed as Gene length corrected trimmed mean of M-values (GeTMM). Right: Scatter plot analogous to Fig. 7C depicting the distribution of Rhino (I) or Kipferl (J) dependent elements in respect to their occupancy by Rhino in MTD-Gal4 > w-sh ovaries, as well as their Rhino-independent Kipferl affinity.

